# A resource-efficient tool for mixed model association analysis of large-scale data

**DOI:** 10.1101/598110

**Authors:** Longda Jiang, Zhili Zheng, Ting Qi, Kathryn E. Kemper, Naomi R. Wray, Peter M. Visscher, Jian Yang

**Affiliations:** Institute for Molecular Bioscience, The University of Queensland, Brisbane, Queensland 4072, Australia; Institute for Advanced Research, Wenzhou Medical University, Wenzhou, Zhejiang 325027, China; Queensland Brain Institute, The University of Queensland, Brisbane, Queensland 4072, Australia

**Author notes:** Correspondence: Jian Yang. These authors contributed equally to this work (Longda Jiang, Zhili Zheng).

## Abstract

The genome-wide association study (GWAS) has been widely used as an experimental design to detect associations between genetic variants and a phenotype. Two major confounding factors, population stratification and relatedness, could potentially lead to inflated GWAS test-statistics and thereby spurious associations. Mixed linear model (MLM)-based approaches can be used to account for sample structure. However, genome-wide association (GWA) analyses in biobank samples such as the UK Biobank (UKB) often exceed the capability of most existing MLM-based tools especially if the number of traits is large. Here, we developed an MLM-based tool (called fastGWA) that controls for population stratification by principal components and relatedness by a sparse genetic relationship matrix for GWA analyses of biobank-scale data. We demonstrated by extensive simulations that fastGWA is reliable, robust and highly resource-efficient. We then applied fastGWA to 2,173 traits on 456,422 array-genotyped and imputed individuals and 2,048 traits on 46,191 whole-exome-sequenced individuals in the UKB.

## INTRODUCTION

The genome-wide association study (GWAS) is a powerful experimental design to detect genetic variants associated with a phenotype of interest. Over the past decade, a number of statistical methods have been developed for GWAS, facilitating the discovery of thousands of genetic loci associated with complex traits and diseases^1,2^. In the early GWAS era, the most commonly used approach was linear or logistic regression^3-11^, which is also the basis of most GWAS software tools^12-14^. The statistical power of a GWAS depends on the proportion of phenotypic variance explained by a variant and the sample size^15^. In other words, for the detection of variants with small effects, a large sample size is required. This can be achieved by a meta-analysis of a large number of cohorts even if the sample size of each individual cohort is limited (e.g. GIANT^16^ and PGC^17^). Due to the substantial decrease in genotyping costs in recent years, sample sizes of GWAS have dramatically increased to 100,000s in single cohorts, such as the UK Biobank (UKB)^18^ and the Biobank Japan Project^19^. These large cohorts not only provide new opportunities to make novel discoveries but also bring challenges in computing especially for methods based on multivariate models. New software tools based on linear regression (LR) have also been developed to accommodate the increasing scale of data, including PLINK2^20^ and BGENIE^18^. Population stratification^21,22^ and relatedness^23,24^ are the two major confounders in GWAS, which could potentially lead to spurious associations if not well-controlled for. In an LR analysis, the effect of population stratification is usually accounted for by fitting the first few eigenvectors (also called PCs) from a principal component analysis (PCA) of the SNP genotypes^25^; the confounding due to relatedness can be avoided by excluding one member of each pair of related individuals based on pedigree or SNP-derived relatedness^14,26^, which, however, results in a loss of power, especially because the proportion of individuals with close relatives in the sample is expected to increase as biobanks get larger^18^.

The mixed linear model (MLM) approach has been widely used in GWAS to control for population stratification and relatedness^27-40^. The basic principle is to test for association between each genetic variant and the phenotype, conditioning on the sample structure inferred from all the genome-wide SNPs^39^. However, the runtime of most existing MLM-based methods ranges from O(*MN*^*2*^) to O(*M*^*2*^*N*)^29,32,34,37-39^, where *M* is the number of variants and *N* is the sample size. Several recent studies have focused on the application of MLM-based methods in biobank-scale data^41-43^. Yet it is still resource demanding to run MLM-based GWAS analyses with millions of genetic variants especially when the number of phenotypes to be analysed is large.

In this study, we propose an extremely resource-efficient approach to perform an MLM-based genome-wide association (GWA) analysis (called fastGWA), implemented in the software GCTA package^26^. We show by extensive simulations that fastGWA is robust in controlling for false positive associations in the presence of population stratification and relatedness, and that fastGWA is ∼89 times faster and only requires ∼5% of RAM compared to the most efficient existing MLM-based GWAS tool in a very large sample (400,000 individuals and 8,531,416 variants). We then demonstrate the utility of fastGWA by analysing GWAS data of 456,422 array-genotyped and imputed individuals (including close relatives) of European ancestry for 2,173 traits and a subset of 49,960 whole-exome sequence-based individuals for 2,048 traits in the UKB. All the summary statistics are publicly available at our data portal, with an online tool to query the association results by genetic variant or phenotype (**URLs**).

## RESULTS

### Overview of the methods

The fastGWA model can be written as

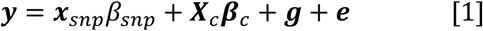

where *y* is an *n* × 1 vector of mean centred phenotypes with *n* being the sample size; *x*_*snp*_ is a vector of mean-centred genotype variables of a variant of interest with its effect *β*_*snp*_; ***X***_*c*_ is the incidence matrix of fixed covariates (e.g., sex, age and the first few PCs) with their corresponding coefficients ***β***_*c*_; ***g*** is a vector of the total genetic effects captured by pedigree relatedness with 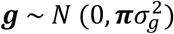; ***π*** is the family relatedness matrix (FAM) based on pedigree structure^44^, e.g., 0.5 for a full-sib or parent-offspring pair; ***e*** is a vector of residuals with 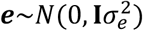. The variance-covariance matrix of *y* is 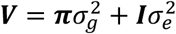. In practice, if pedigree information is missing or largely incomplete, ***π*** can be replaced by an SNP-derived genetic relationship matrix (GRM) with all the small off-diagonal elements (e.g., those < 0.05) set to zero. This is because it has been shown in a previous study^45^ that the sparse GRM thresholded at 0.05 captures approximately the same proportion of phenotypic variance as the FAM does (confirmed by our simulation below). Here we present two closely related versions of our method, the fastGWA, based on sparse GRM computed from genotype data, and the fastGWA-Ped, based on FAM constructed from pedigree information.

The fastGWA model imposes control over relatedness by pedigree information or realised sparse GRM with the effect of population stratification captured by the SNP-derived PCs. The variance components 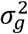 and 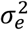 are unknown but can be estimated by the restricted maximum likelihood (REML) algorithm^46^. We have implemented in fastGWA an efficient grid-search-based REML algorithm (termed as fastGWA-REML) to estimate 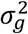 and 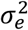 without the need to compute ***V***^-1^ (**Supplementary Note 1**). In the presence of moderate to strong common environmental effects shared among relatives, the genetic variance estimated from closely related individuals (e.g., pairs of individuals with relatedness coefficients > 0.05, see **Methods**) may be a better quantity than that estimated based on genetic relatedness between all pairwise individuals in the sample as in most existing MLM-based methods^27-40^. This is because 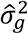 estimated from close relatives captures the variation due to the overall additive genetic effect as well as that due to common environmental effects (see **Discussion** for details).

Once the estimates of variance components are obtained, the variance-covariance matrix ***V*** and its inverse can be computed efficiently using the sparse matrix algorithms implemented in Eigen C++ library (**URLs**). Therefore, *β*_*snp*_ can be estimated using the generalised least squares approach:

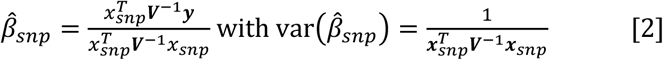

where the parameters used to compute ***V*** are unknown but can be replicated by the estimates from sparse REML under the null that *β*_*snp*_ = 0 as in most existing methods^27,29-31,37,39,40^. The computational efficiency of the association test can be improved by orders of magnitude using the GRAMMAR-GAMMA approximation^36^ (**Supplementary Note 2**). We have implemented fastGWA in the GCTA software^26^ with a user-friendly command-line interface (**URLs**). We limited all the analyses in this study to common variants because of the known limitations of MLMs in rare-variant association^42,47^.

### Runtime and resource requirements

Given that fastGWA is specifically designed for large-scale data, we chose to evaluate its computational performance (i.e., runtime and resource requirements) using the UKB^18^ data (456,422 individuals of European ancestry and 8,531,416 variants with MAF ≥ 0.01) (**Methods**). We confirmed by simulations that the estimate of 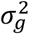 from the fastGWA-REML algorithm was nearly identical to that from the average information^48^ REML algorithm implemented in GCTA^26^ (**Supplementary Figure 1**) and that the fastGWA test-statistic computed using the GRAMMAR-GAMMA approximation was almost the same as that using the exact approach (**Supplementary Figure 2**). We randomly sampled subsets of individuals from the UKB (*n* = 50,000 to 400,000) and compared fastGWA in GCTA v1.92.3 with the infinitesimal mixed model in BOLT-LMM v2.3.2^40,42^ and the linear regression (LR) model in PLINK2 v2.00a2 on a computing platform with 96 GB memory and 16 CPU cores with the runtime capped at 7 days (168 hours). The tests were performed using a real trait, body mass index (BMI), with an estimated SNP-based heritability of ∼0.27^49,50^. The genotype data were stored in PLINK binary PED format^14,20^. Each test was repeated 10 times to obtain an average. The results showed that GCTA-fastGWA completed the analysis in ∼20 minutes for the 400K sample – approximately 1.1% of that of BOLT-LMM (∼22 hours) (**Table 1**). While BOLT-LMM requires a running time of *O*(*MN*^1.5^), the time complexity of fastGWA is approximately *O*(*MN*), almost the same as that of LR. GCTA-fastGWA was even ∼4 times faster than PLINK2, likely because of coding efficiency and the difference in strategy used to deal with missing genotypes (note that the LR version of fastGWA was also 4 to 5 times faster than PLINK2). A detailed speed comparison of the three methods can be found in **Table 1** (tested on solid-state drives) and **Supplementary Table 1** (tested on hard disk drives). BOLT-LMM is optimized for genotype data in BGEN v1.2 format^51^ (Po-Ru Loh, personal communication) and our benchmark testing confirmed that BOLT-LMM with data in BGEN format reduced the runtime of the “association” procedure (**Table 1**) by approximately ½ but did not have significant improvement on the other procedures. While all the tests of the three methods were conducted on the same computing platform (i.e., 96 GB memory and 16 CPU cores), the actual memory usage differed substantially. GCTA-fastGWA used much less resource than BOLT-LMM (**Table 2**). For a data set with sample size of 400K, GCTA-fastGWA required less than 5 GB of memory to complete the whole computation, only ∼5% of the usage of BOLT-LMM. Note that we did not report the memory usage of PLINK2 as it preserved as much memory as possible according to the overall physical memory. If the pedigree information is not available, we may need to take the computing cost of the GRM into consideration for GCTA-fastGWA (**Supplementary Note 3**) and PLINK2-LR (because of the use of GRM to exclude related individuals in practice although the LR analyses above were performed using all individuals for a fair comparison). Nevertheless, the GRM computation is often part of the quality control (QC) process and only needs to be done once for the analyses of all traits, meaning that the additional computational cost per trait on average for GCTA-fastGWA or PLINK2-LR due to GRM computation is not very expensive. In addition, we showed that the runtime of fastGWA-REML was approximately linear to sparse GRM density (**Supplementary Table 2**). Since the variance estimation step only needs to be done once under the null model and the time complexity of the association test step is nearly independent of sparse GRM density (**Supplementary Note 2**), the effect of sparse GRM density on the overall runtime of fastGWA is limited in a certain range.

**Table 1.** Comparison of runtime between fastGWA, BOLT-LMM, and PLINK2 Shown are the runtimes of GCTA-fastGWA v1.92.3 (mixed model) and BOLT-LMM v2.3.2 (infinitesimal mixed model), and PLINK2 v2.00a2 (LR model). The data used in this test consisted of 8,531,416 common variants, of which 565,631 LD-pruned variants were used as “model SNPs” in BOLT-LMM (**Supplementary Note 3**). The runtime of fastGWA or BOLT-LMM consists of two steps: a) the estimation of mixed model parameters (“Para. Est.”), and b) the association test (“Assoc.”). The LR was performed using all the individuals. All tests were performed the same computing environment: 96 GB memory and 16 CPU cores with solid-state disk in one computer node. The computing time of the GRM (as required by fastGWA and LR) is described in **Supplementary Note 3**.

| Sample Size | GCTA-fastGWA |  |  | BOLT-LMM |  |  | PLINK2 |
| --- | --- | --- | --- | --- | --- | --- | --- |
|  | Para. Est.<br>(h) | Assoc.<br>(h) | Total<br>(h) | Para. Est.<br>(h) | Assoc.<br>(h) | Total<br>(h) | Total<br>(h) |
| 50,000 | 0.00 | 0.03 | 0.03 | 0.88 | 1.05 | 1.93 | 0.07 |
| 100,000 | 0.00 | 0.04 | 0.04 | 2.09 | 2.07 | 4.16 | 0.15 |
| 200,000 | 0.01 | 0.07 | 0.08 | 5.34 | 4.16 | 9.50 | 0.37 |
| 300,000 | 0.01 | 0.14 | 0.15 | 9.51 | 6.24 | 15.75 | 0.81 |
| 400,000 | 0.02 | 0.23 | 0.25 | 13.85 | 8.44 | 22.29 | 1.15 |

**Table 2.** Comparison of memory usage between fastGWA and BOLT-LMM Shown are the actual memory (“Mem”) and virtual memory (“VMem”) usages of GCTA-fastGWA v1.92.3 and BOLT-LMM v2.3.2 (infinitesimal model only) in GB units. The data used in this test consisted of 8,531,416 common variants, of which 565,631 LD-pruned variants were used as “model SNPs” in BOLT-LMM (**Supplementary Note 4**). All tests were performed in the same computing environment: 96 GB memory and 16 CPU cores in one computer node.

| Sample size | GCTA-fastGWA | | BOLT-LMM | | $\frac{Mem_{fastGWA}}{Mem_{BOLT-LMM}}$ | $\frac{VMem_{fastGWA}}{VMem_{BOLT-LMM}}$ |
| --- | --- | --- | --- | --- | --- | --- |
|  | Mem (GB) | VMem (GB) | Mem (GB) | VMem (GB) |  |  |
| 50,000 | 1.4 | 3.3 | 8.8 | 9.7 | 15.9% | 34.0% |
| 100,000 | 1.5 | 3.3 | 15.3 | 16.2 | 10.5% | 20.4% |
| 200,000 | 1.8 | 3.3 | 28.6 | 29.6 | 6.3% | 11.5% |
| 300,000 | 2.2 | 4.3 | 42.0 | 43.1 | 5.2% | 10.0% |
| 400,000 | 2.7 | 4.7 | 55.5 | 56.6 | 4.9% | 8.3% |

**Figure 1.**
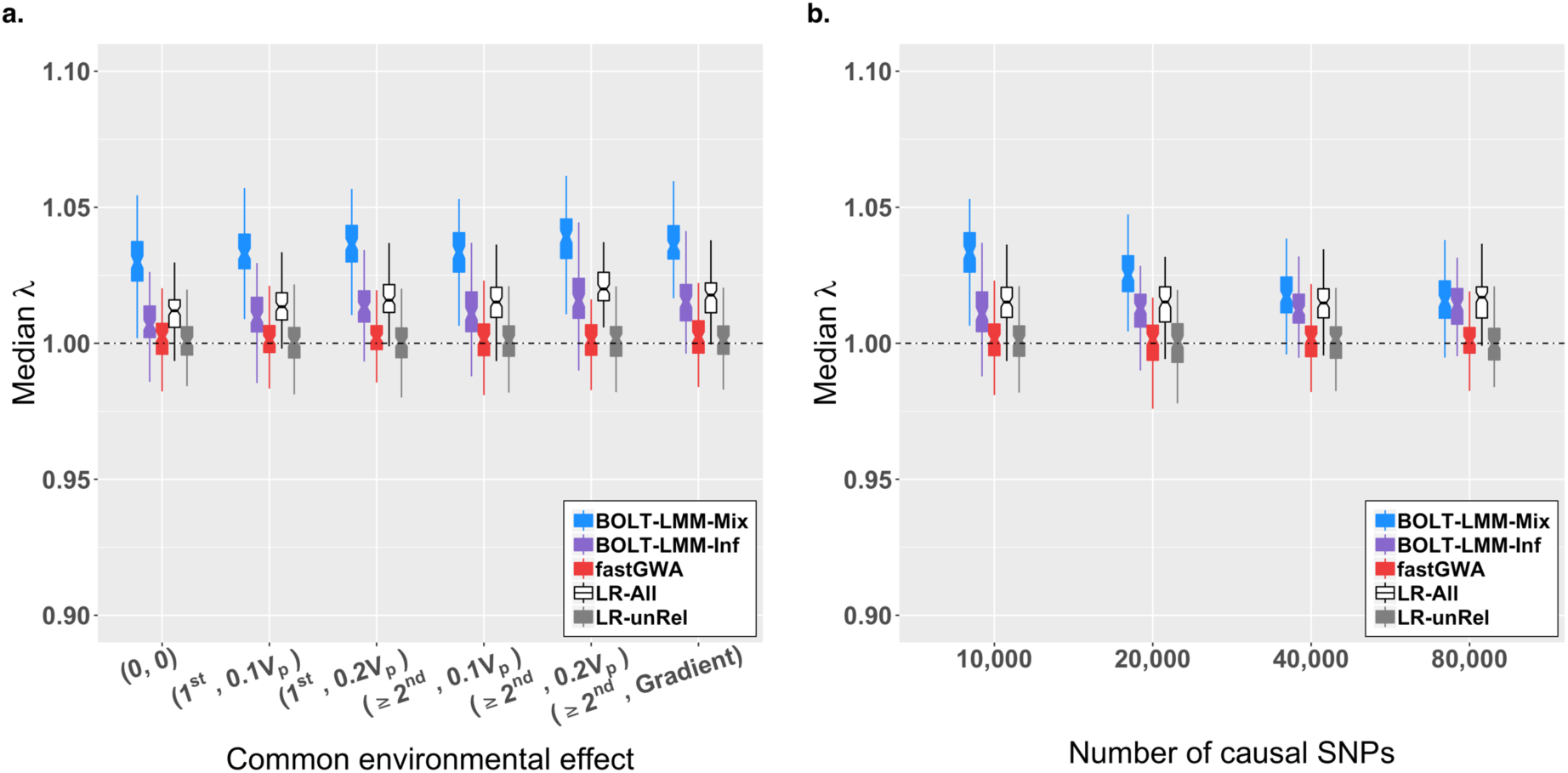
Median *λ* of null variants under different simulation scenarios. a) Median *λ* of null variants with different levels of common environmental effects in the simulations (**Methods**). The y axis represents the median *λ* of the null variants (i.e., all the variants on the even chromosomes), and the x axis represents different levels of common environment effects with (0, 0) representing no common environmental effects; (1^st^, 0.1V_p_) and (1^st^, 0.2V_p_) representing common environmental effects explaining 10% and 20% of the phenotypic variance (V_p_) among the 1^st^ degree relatives, respectively; (≥2^nd^, 0.1V_p_) and (≥2^nd^, 0.2V_p_) representing common environmental effects explaining 10% and 20% of V_p_ among the 1^st^ and 2^nd^ degree relatives (note that this actually includes all the relatives simulated), respectively; (≥2^nd^, Gradient) representing 20% of V_p_ among the 1^st^ degree relatives and 10% of V_p_ among the 2^nd^ degree relatives. Each boxplot represents the distribution of median *λ* across 100 simulation replicates. b) Median *λ* of null variants with different number of causal variants (i.e., 10k, 40k, and 80k)) in the simulations (**Methods**). The y-axis represents the median *λ* of the null variants and the x-axis represents the different number of causal variants simulated.

**Figure 2.**
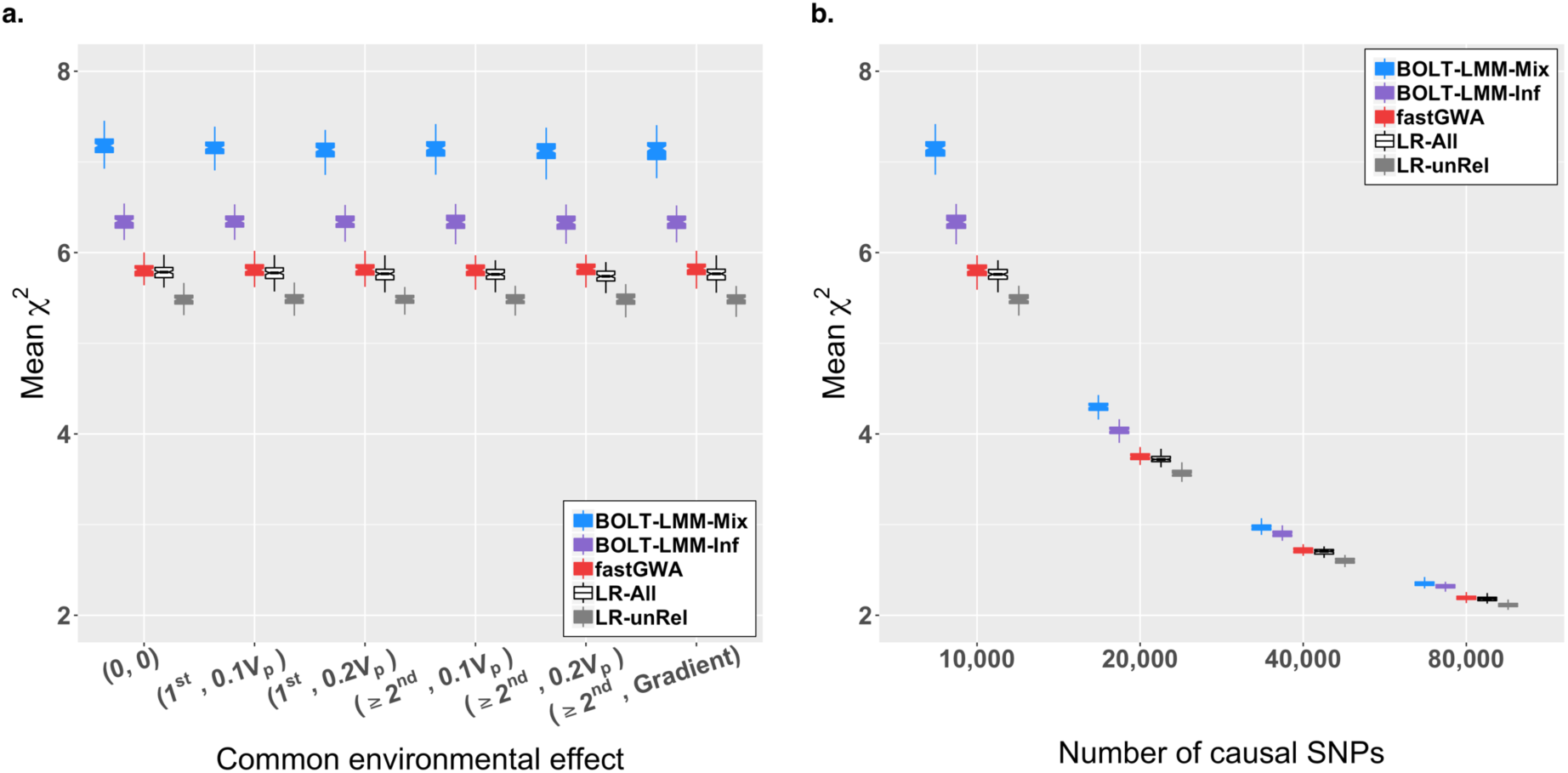
Mean *χ*^2^ of causal variants under different simulation scenarios. a) Mean *χ*^2^ of causal variants with different levels of common environmental effects in the simulations (**Methods**). The y-axis represents the mean *χ*^2^ values for all the 10,000 causal variants on the odd chromosomes and the x-axis represents the different levels of common environmental effects as described in **Figure 1a**. The mean *χ*^2^ has been adjusted by the median *λ* of null variants on the even chromosomes. Each boxplot represents the distribution of mean *χ*^2^ across 100 simulation replicates. b) Mean *χ*^2^ at causal variants with different number of causal variants in the simulations (**Methods**). The y-axis represents the average of mean *χ*^2^ of the causal variants and the x-axis represents the different number of causal variants (i.e., 10k, 20k, 40k, 80k). The mean *χ*^2^ has been adjusted by the median *λ* of null variants on the even chromosomes. Each boxplot represents the distribution of mean *χ*^2^ across 100 simulation replicates.

### False positive rate and statistical power

We used extensive simulations to quantify the genomic inflation factor, the false positive rate (FPR, number of false positive associations divided by the total number of tests) and the statistical power of fastGWA in comparison with LR in PLINK2 and MLMs in BOLT-LMM (**Methods**). A sample of 100,000 individuals was generated by random sampling of chromosome segments from a subset of the UKB data to mimic a cohort with substantial population stratification and relatedness (**Supplementary Notes 4** and **5**; **Supplementary Figures 3-6**). One of the main aims of this simulation study is to investigate the influences of common environmental effects on different association test methods. We generated phenotypes from a number of causal variants randomly sampled from all variants on the odd chromosomes, leaving those on the even chromosomes as the null variants to quantify the genomic inflation/deflation in test-statistics and the FPR. We mimicked the effect of population stratification by generating mean phenotype difference between two populations and the effect of relatedness by specifying different degrees of common environmental effects among close relatives (**Methods**). The genomic inflation factor, or *λ*_GC_, is defined as the median chi-squared statistic divided by its expected value at the null variants^21,52^.

**Figure 3.**
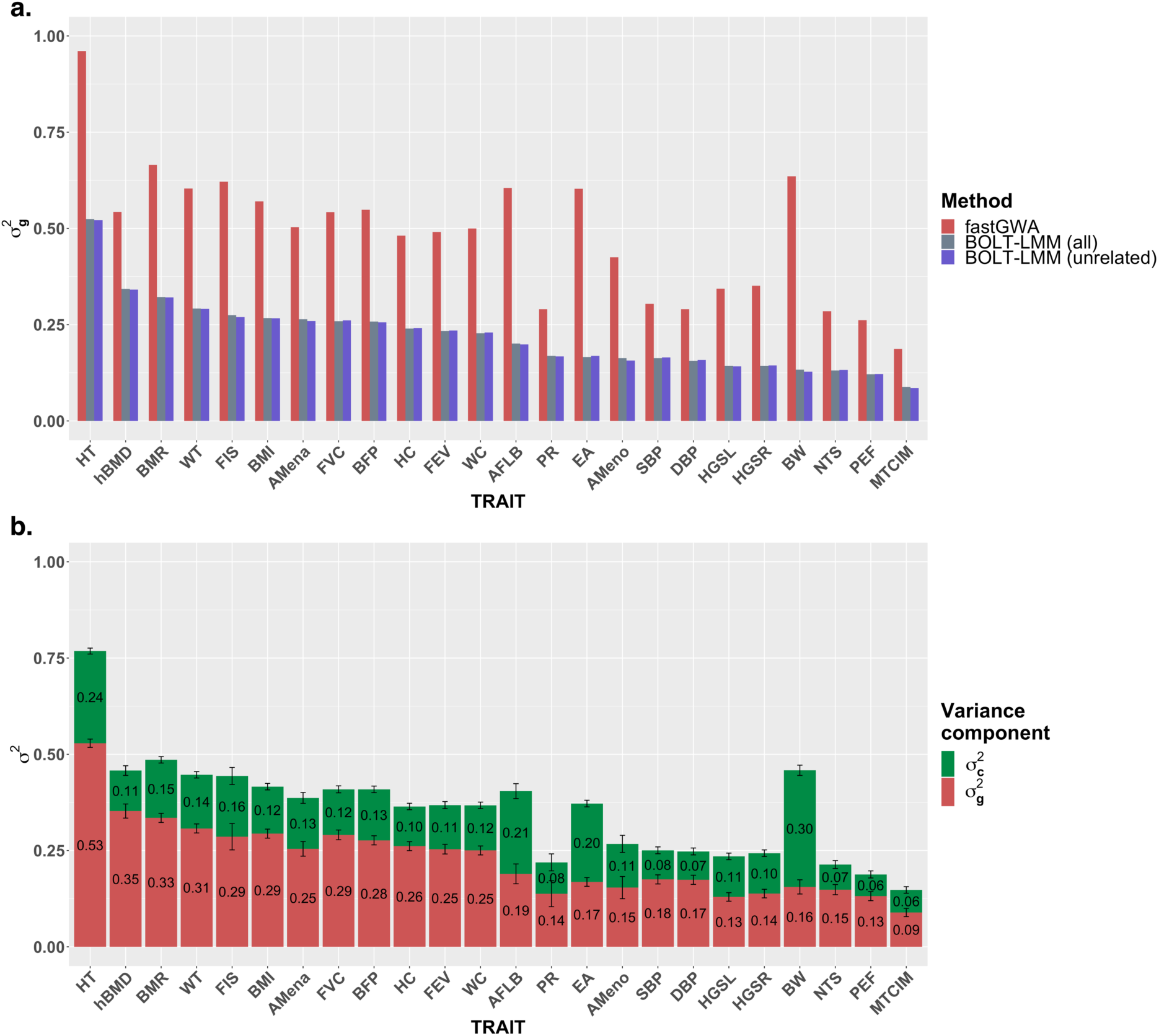
Estimates of genetic variance by fastGWA and BOLT-LMM for 24 traits in the UKB. A full list of phenotype abbreviations can be found in **Supplementary Table 3**. Shown in panel a) are the estimates of genetic variance (i.e., 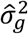) by fastGWA-REML using all individuals, by BOLT-REML^40,42^ (used in BOLT-LMM) using all individuals, and by BOLT-REML^59^ using unrelated individuals. In panel b), we analysed a subset of the UKB data (21,815 inferred full-sib pairs, comprising 39,934 individuals from 19,386 families; **Supplementary Note 3**) based on a two-component model: *y* = ***g*** + ***e***_***c***_ + ***e*** with 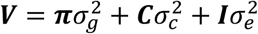 where ***g*** is a vector of total genetic effects, ***e***_***c***_ is a vector of shared environmental effects, ***π*** is the full dense GRM estimated from a set of slightly-pruned HapMap3 SNPs as used in real data analyses (*m* = 565,631, see **Supplementary Note 3** for details), and ***C*** is a design matrix with 1 or 0 to indicate whether a pair of individuals belong to the same family. Standard errors of the estimates are represented by the error bars.

Our simulation results showed that there was inflation in the test-statistics of null variants from LR-All (i.e., LR analysis of all the individuals including close relatives) even in the absence of common environmental effects (**Figure 1a**). This is because of the inter-chromosome correlations between the causal and null variants induced by the relatedness in the sample. The test-statistics of null variants from BOLT-LMM-Mix (a Bayesian mixture model) were inflated, and the inflation was higher than that of LR-All. BOLT-LMM-Inf (BOLT-LMM infinitesimal model), which is a special case of BOLT-LMM-Mix, assumes only a single distribution of the variant effects. Note that BOLT-LMM-Inf is similar but computationally more efficient than the MLM leave-one-chromosome-out (LOCO)^39^ approach implemented in GCTA. We also observed inflation in the test-statistics of null variants from BOLT-LMM-Inf and the inflation increased as the variance explained by common environmental effects increased (**Figure 1a**), likely because BOLT-LMM-Inf failed to capture the variance due to common environmental effects by fitting all common variants on the other chromosomes as random effects (**Supplementary Figure 7**; see below for more discussion).

In contrast, there was almost no inflation for both LR-unRel (i.e., LR analysis restricted to unrelated individuals) and fastGWA. As the level of common environmental effects increased, the genomic inflation factors of LR-All, BOLT-LMM-Mix, and BOLT-LMM-Inf all increased slightly, but not for LR-unRel and fastGWA, demonstrating the robustness of fastGWA in accounting for common environmental effects shared among relatives. We also found that the results from fastGWA-Ped were very similar to those from fastGWA (**Supplementary Figure 8**, recognising that pedigree relationships were known without error in simulation). Additionally, we quantified the FPR using the null variants (i.e., all the variants on the even chromosomes), where FPR is defined as the proportion of null variants with p-values < 0.05 in each simulation replicate. The FPRs of the methods were in line with their observed genomic inflation factors (**Supplementary Figure 9**).

Next, we extended the simulation with larger numbers of causal variants. We kept the proportions of variance explained by common environmental effects and population stratification the same as that in one of the scenarios above (i.e., common environmental effects explained 10% of phenotypic variance (*V*_*p*_) among all relatives and population stratification explained 5% of *V*_*p*_). The results were similar to those presented above, i.e., the test-statistics were inflated for BOLT-LMM and LR-All but not for fastGWA and LR-unRel (**Figure 1b**). The inflation in test-statistics from BOLT-LMM-Mix decreased as the number of causal variants increased (**Figure 1b**). It is of note that, in all the simulation scenarios where there were shared environmental effects, the test-statistics at the null variants from BOLT-LMM-Mix appeared to be even more inflated than those from LR-All (i.e., linear regression without correcting for relatedness).

To quantify the statistical power of each method, we used the mean *χ*^2^ statistic at the causal variants. Because the test-statistics from some of the methods were inflated at the null variants, we divided the mean *χ*^2^ statistic at the causal variants by the genomic inflation factor at the null variants to compare the power of different methods given the same level of FPR (**Methods**), similar to the idea of computing the area under the power-FPR ROC curve. We found that BOLT-LMM-Mix showed the highest power among all the methods (**Figure 2a**). When the number of causal variants was relatively small, there was a relatively large gap in power between BOLT-LMM-Mix and all other methods including BOLT-LMM-Inf (**Figure 2a**). BOLT-LMM-Inf model showed the second highest power, in line with the theory that MLM leaving out the target chromosome from the polygenic component gains power^39^. fastGWA showed a similar level of power to LR-All, and LR-unRel showed the lowest power among all the methods owing to its smaller sample size. We also observed that the power of all the methods were almost independent of the variance explained by common environmental effects. Increasing the number of causal variants led to smaller differences in power between methods (**Figure 2b**), suggesting that the difference in power increased as the per-variant variance explained increased (**Methods**).

We further assessed the robustness of fastGWA by simulation in a few additional scenarios. The results show that the test-statistics of fastGWA remained to be well calibrated for case-control phenotypes regardless whether cases were oversampled (**Supplementary Figures 10** and **11**), for non-normally distributed phenotypes (**Supplementary Figure 12**), or in a sample with substantially higher sparse GRM density than that in the UKB (**Supplementary Figures 13** and **14**).

### Application of fastGWA to 2,173 traits in the UKB

We used fastGWA to conduct a genome-wide association analysis of 8,531,416 variants with MAF ≥ 0.01 in all the UKB individuals of European ancestry (*n* = 456,422) for 2,173 real phenotypes and compared the results to those produced by the Neale Lab using LR-unRel (361,194 unrelated individuals, see **URLs**). As noted above, fastGWA is expected to be more powerful than LR-unRel because of the larger sample size. We followed the QC and analysis pipelines as used in the Neale Lab’s analysis with some modifications (e.g., we kept all the related individuals) (**Methods**). We confirmed that the test statistics were highly correlated between the two sets of results (mean correlation of z-statistics of 0.89 for 1,163 overlapping phenotypes). We chose 24 representative traits (**Supplementary Table 3**) to compare our results with the Neale Lab’s results.

We first attempted to quantify the genomic inflation in the two sets of results. LD score regression (LDSR) is an approach developed to partition the inflation in GWAS test-statistics into components due to polygenic variation and population structure^53^, but recent studies suggest that LDSR intercept is a function of heritability, sample size and population genetic differentiation in the sample, and is expected to deviate from 1 in a genetically stratified sample even if the phenotype is not stratified^42,53,54^. We therefore sought to quantify the inflation due to sample structure by the attenuation ratio, i.e., (LDSC intercept – 1) / (mean *χ*^2^ − 1), which is independent of sample size, as suggested in a recent study^42^. The attenuation ratio of fastGWA was very close to that of LR-unRel on average across the 24 traits (0.0792 vs. 0.0783; **Supplementary Table 4**), consistent with what we observed in simulations (**Figure 1**) that the inflation due to relatedness can be reasonably well corrected for by fastGWA.

We then compared the discovery power between the two sets by the clumping analysis in PLINK2^20^ (*P*-value threshold = 5×10^−9^, window size = 5 Mb, and LD *r*^2^ threshold = 0.01). Of all the 24 traits, the number of clumped genome-wide significant variants was 7,839 in our results, substantially higher than that (5,676) in the Neale Lab’s results (see **Supplementary Table 5** for the comparison of each trait), suggesting a nearly 40% of improvement in the number of GWAS discoveries for fastGWA over LR-unRel. In addition, Canela-Xandri et al.^55^ have also applied an MLM-LOCO^39^ association analysis to 778 UKB traits by DISSECT^41^ using parallel computing in a supercomputer and released all the GWAS summary data in a public database, GeneATLAS (**URLs**). We compared the results from GeneATLAS to those from the Neale Lab and our fastGWA analysis for 10 traits available in all the three sets (**Supplementary Table 4** and **5**). There was no significant difference in attenuation ratio between the three sets but GeneATLAS had more clumped variants than the other two sets, likely because of the MLM-LOCO scheme used in GeneATLAS^55^, consistent with the simulation results from this study (**Figure 2**) and previous studies^39,40^ that MLM-LOCO gains power. We also analysed the 24 traits using BOLT-LMM-Inf which showed a larger number of clumped variants (**Supplementary Table 4**) but with a slightly higher attenuation ratio compared to any of the other three methods (**Supplementary Table 3**).

During the revision of this manuscript, whole-exome sequence (WES) data of 49,960 participants became available in the UKB. We therefore applied fastGWA to the WES data (152,327 variants with MAF ≥ 0.01 and missingness rate ≤ 0.1, and 46,191 individuals of European descent) for 2,048 traits, following the same analysis pipeline described above (**Methods**). We identified 148 near-independent associations at an exome-wide significance level (*P*-value < 0.05/ *m* with *m* being the number of variants tested for a trait) for the 24 traits described above (**Supplementary Table 6**). For each of the exome-wide significant associations, we repeated the fastGWA analysis conditioning on the GWAS signals (within 10Mb of the WES variant in either direction) identified from the imputed data of the whole UKB sample described above. Conditioning on the imputed GWAS signals, only 4 associations remained exome-wide significant (**Supplementary Table 7**), suggesting that most common variants in the WES data have been well tagged by SNP array-based genotyping and imputation. Full summary statistics of 8,531,416 array-genotyped or imputed variants for 2,173 traits and 152,369 WES variants for 2,048 traits are publicly available at our data portal without restricted access (**URLs**). Additionally, we developed an online tool for users to query and visualize the UKB summary statistics (**URLs**).

## DISCUSSION

In this study, we developed a reliable, robust and resource-efficient association analysis tool, fastGWA, which requires much smaller system resources (i.e., runtime and memory usage) than existing tools, making it feasible to conduct GWA analyses of thousands of traits in large cohorts like the UKB without the need to remove related individuals. The tool is also applicable to family-structured data with a very large number of molecular phenotypes.

Apart from computational efficiency, fastGWA also shows greater robustness than existing MLM-based methods in the presence of confounding factors. It has long been known that the existence of relatedness in the data would lead to inflated association test statistics^23,24,56^, confirmed by our simulation (**Figure 1a**). MLMs can be used to account for relatedness because the fixed effect is tested conditional on the phenotypic covariance structure among all individuals (**Equation 1**)^44^. It should be noted that the estimate of the “genetic variance” component based on close relatives as in fastGWA is a compound of *V*_*g*_ (the true genetic variance) and *V*_*C*_ (the amount of phenotypic variance attributable to shared environmental effect). More specifically, in our simulated data, the phenotypic covariance between two close relatives is *cov*(*y*_*i*_, *y*_*j*_)= *π*_*ij*_*V*_*g*_ + *V*_*C*_ with *π*_*ij*_ being the family relatedness. In the fastGWA analysis, however, we do not seek to explicitly partition *V*_*g*_ and *V*_*C*_ but to use a single variance component to model the phenotypic covariance among close relatives. In the fastGWA model, the deviation of 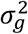 (being estimated based on the sparse GRM) from *V*_*g*_ is a function of 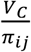 and the proportion of related pairs in the sample. In contrast, the estimate of the “genetic variance” component from an MLM analysis based on a dense GRM (SNP-derived GRM between all pairwise individuals, as in BOLT-LMM) is a weighted average of the SNP-based genetic variance (often smaller than *V*_*g*_ in reality because of imperfect tagging) and the pedigree-based genetic variance (similar to 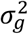 in the fastGWA model), leading to an under-estimation of the covariance between closely related individuals, especially when the proportion of relative pairs is very small compared to the unrelated pairs. It was shown in our simulations that when *V*_*C*_ = 0, the estimate of the “genetic variance” component from either fastGWA or BOLT-LMM was unbiased (**Supplementary Figure 7**). As *V*_*C*_ increased, the estimate of the “genetic variance” component from fastGWA increased but the estimate from BOLT-LMM-Inf was almost unchanged (**Supplementary Figure 7**). This may explain the inflation of test-statistics in BOLT-LMM-Inf at the null variants observed in our simulations with non-zero *V*_*C*_ (**Figure 1**). These observations are in line with the simulation results from a recent study^57^. To further demonstrate the issue above, we selected 24 real phenotypes from the UKB to estimate the genetic variance using different methods (the same 24 traits as used in the UKB real data analyses except for education attainment which was reconstructed following the method in Ref.^58^). Our result shows that the BOLT-REML^59^ (the method used in BOLT-LMM to estimate the variance components) estimate of “genetic variance” is equivalent to the estimate of genetic variance corresponding to the full dense GRM (**Figure 3**), leaving the covariance between close relatives due to common environmental effect (and/or rare variants) uncaptured (**Figure 3**). Two particular examples were educational attainment (EA) and birth weight (BW), which have been shown in previous studies with strong common environmental effects (e.g., shared maternal effect among sibs)^60-62^. The estimate of the “genetic variance” component for EA and BW from fastGWA were much higher than those from BOLT-REML (**Figure 3**), consistent with a substantial estimate of 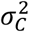 in a two-component model (**Figure 3**). All these results caution the use of BOLT-LMM for traits with large *V*_*C*_.

The increased power of fastGWA compared to LR-unRel is mainly because more individuals are included in the association test. Large population-based cohorts such as the UKB tend to oversample relatives as participants when an assessment-centre based recruitment strategy is implemented^18^. Taking the UKB cohort as an example, to generate a set of unrelated individuals, 107,864 out of 456,422 European participants need to be excluded given a relatedness threshold of 0.05 in GCTA. Excluding these participants would significantly compromise the statistical power, which can be avoided by implementing MLM-based methods such as fastGWA. In addition, the higher power of BOLT-LMM compared to fastGWA or LR is mainly driven by its LOCO scheme. Previous studies have showed that leaving the target chromosome out of the polygenic component gains power because the effects of variants on the other chromosomes are conditioned out from the model and proximal contamination (i.e., the target variant being fitted twice in the model, once as a fixed effect and again as a random effect) is avoided^32,39,40^. We did not observe any increase in power when applying the LOCO scheme to fastGWA (**Supplementary Figure 15**) because fastGWA estimates pedigree relatedness by a sparse GRM to model phenotypic covariance between close relatives due to genetic and/or common environmental effects and the pedigree relatedness estimated using all autosomes are similar to those using 21 chromosomes under the LOCO scheme. It is of note that to save computational time, BOLT-LMM-Inf estimates the genetic variance only once using all “model SNPs” and applies it to the association tests of all variants without re-estimating the genetic variance when a chromosome is left out, assuming that genetic variance attributable to a single chromosome is relatively small. This approximation could give rise to inflated test-statistics of BOLT-LMM-Inf under the null even in the absence of shared environmental effects^40^ (see the “(0,0)” scenario in **Figure 1**). This issue could be fixed by re-estimating the variance components when a chromosome was left out from the polygenic component (**Supplementary Figure 16**). In addition, BOLT-LMM-Mix requires LD scores to calibrate the test statistics^40^. We observed from simulations that there was an effect on the choice of LD reference on the BOTL-LMM-Mix test-statistics (**Supplementary Figure 17**), which may explain part of the inflation in BOLT-LMM-Mix test-statistics under the null (**Figure 1**).

There are several caveats of applying fastGWA in practice. First, if the pedigree information is unavailable or incomplete (as is the case for UKB; shown in **Supplementary Figure 18** and further discussed in **Supplementary Note 3**), it is necessary to compute the GRM from SNP data. We have implemented in GCTA a very efficient tool to compute the SNP-based GRM along with a function that can subdivide the GRM computation into a large number of components for parallelized computing (**Supplementary Note 3**). These GRM components can be finally assembled to a full GRM using a simple but efficient Linux/Unix command. Second, fastGWA uses SNP-derived PCs to correct for the effect due to population stratification. PCs are often provided as part of the QC package in the downloaded data^18^. If PCs are not available, we would recommend efficient PCA tools such as fastPCA or FlashPCA^63,64^. Another more efficient approach is to compute PCs in a subset of the sample, and project the PCs to the rest of the sample. This approach has been implemented in GCTA (**URLs**). It is likely that PCs are also required for other MLM-based methods including BOLT-LMM because although in theory MLM-based methods accounts for population stratification by fitting all (or a subset of selected) variants as random effects^29,30,39^, MLM-based association analyses in large samples suggest that fitting PCs as covariates improves robustness^42,43^. It is also noteworthy that computing PCs from all the variants might be suboptimal, as the test-statistics would tend to be deflated under the null (**Supplementary Note 6** and **Supplementary Figure 19**). It is therefore recommended to compute PCs from a set of LD-pruned variants. Third, as discussed above, in the presence of common environmental effects, “genetic variance” component in the fastGWA model is a function of *V*_*g*_ and *V*_*C*_. We did not attempt to differentiate common environmental components among different degrees of relatedness (e.g., siblings might share stronger common environmental effects than cousins). Nevertheless, our simulation showed that this simple modelling did not lead to inflated test-statistics at the null variants in the scenario where common environmental effects decreased as relatedness decreased (**Figure 1**). Fourth, the fastGWA program will switch to use LR for analysis (allowing for covariates) if the estimate of the genetic variance component is not significant at a nominal significance level, cautioning the use of fastGWA in a sample with a small number of related pairs. Last but not least, this study is focused on common variants, and fastGWA likely suffers from the same weakness as the other MLM-based methods in rare variant association test^47^ especially for skewedly distributed phenotypes or binary traits with very low prevalence rate^42^.

Despite these caveats, fastGWA is an MLM-based association analysis method that is orders of magnitude more resource-efficient and has more robust control over relatedness than existing methods. The computational efficiency of fastGWA has been manifested by its successful application to the genome-wide association analyses of 2,173 traits on 456,422 array-genotyped in the UKB. The summary statistics released from this study are useful resources for post-GWAS analyses (e.g., functional enrichment, genetic correlation, polygenic risk score, and causal inference) and phenome-wide association studies (PhWAS). Additionally, fastGWA can be modified for omic-data-based QTL (xQTL) analyses in biobank samples in the future.

## METHODS

### UK Biobank data

The UK Biobank (UKB) is a large cohort study consists of approximately half a million participants aged between 40 and 69 at recruitment, with extensive phenotypic records^18^. In this study, we selected 456,422 individuals of European ancestry from the UKB cohort for simulation and real data analyses. Genetic data were genotyped by two different arrays, the Applied Biosystems^™^ UK Biobank Axiom^™^ Array and the Applied Biosystems^™^ UK BiLEVE Axiom^™^ Array^18^, of which 556,269 genotyped variants were for simulation and 8,531,416 variants imputed by UKB consortium (imputed Version 3) were used for real data analyses^18^. Genotyped/imputed data were filtered with standard QC criteria in PLINK2^20^, e.g., MAF ≥ 0.01, Hardy-Weinberg Equilibrium test *P* ≥ 10^−6^, genotyping rate ≥ 0.95, and imputation info score ≥ 0.8 in real data analyses. In addition, the UKB released its first tranche of whole-exome sequence (WES) data of 49,960 participants in March 2019^65^. The WES variants had been called and cleaned by two different pipelines, Regeneron’s Seal Point Balinese (SPB)^65^ and Functionally Equivalent (FE)^66^. We used the FE data for analysis and excluded from the analysis variants with MAF < 0.01 and missingness rate > 0.1 (152,327 variants remained) and individuals with non-European ancestry (46,191 individuals remained).

### Simulating genotypes

In order to test the performance of fastGWA in the presence of relatedness and substantial population stratification, we simulated a total of 100,000 artificial individuals from two different ancestry backgrounds, with a moderate proportion of relatives (10% of all samples) using a “mosaic-chromosome” scheme modified from^40^. We first selected all individuals with self-reported “British” and “Irish” ancestry from the UKB as the founders. We then filtered the samples based on their genetic ancestry inferred from SNP data to assure that the two groups were genetically distinct (**Supplementary Note 4** and **Supplementary Figure 3-4**). Next, we divided the genome into consecutive segments of 2,000 variants and generated unrelated individuals by selecting each segment from two of the founders chosen at random, and simulated related individuals by selecting the segments from a limited number of founders according to the relatedness (**Supplementary Note 4**). Finally, we obtained 45,000 independent and 5,000 related “British individuals”, and 45,000 independent and 5,000 related “Irish individuals”. Detailed description of the parameters and procedures have been described in the **Supplementary Note 4**. We used GCTA to compute LD scores from the simulated genotype data (**Supplementary Note 7**).

### Simulating phenotypes

The phenotypes were simulated based on the following model

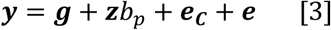

where 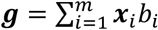, is the sum of the genetic effect of *m* causal variants with *x*_*i*_ a vector of variant genotypes and *b*_*i*_∼*N*(0,1); **z** is a vector consisting of 0 (British) and 1 (Irish) to indicate ancestry with *b*_*p*_ being the mean difference in phenotype between the two groups; ***e***_***c***_ is a vector of shared environmental effects with the individual(s) in each family assigned by the same value generated from *N*(0,1); and ***e*** is a vector of individual environmental effects (i.e., the residuals) with *e*_*j*_∼*N*(0, 1). We considered different levels of relatedness in different simulation scenarios including:

a. no common environmental effects, denoted by (0,0);
b. common environmental effects explaining 10% or 20% of the total phenotypic variance (*V*_*p*_) among the 1^st^ degree relatives, denoted by (1^st^, 0.1*V*_*p*_) and (1^st^, 0.2*V*_*c*_)), respectively;
c. common environmental effects explaining 10% or 20% of *V*_*p*_) among the 1^st^ and 2^nd^ degree relatives, denoted by (≥2^nd^, 0.1*V*_*p*_)) and (≥2^nd^, 0.2*V*_*p*_)), respectively;
d. common environmental effects explaining 20% of *V*_*p*_ among the 1^st^ degree relatives and 10% of *V*_*p*_ among the 2^nd^ degree relatives, denoted by (≥2^nd^, Gradient).

Each simulation was repeated 100 times. Detailed description of the parameter settings and details of extended simulations can be found in the **Supplementary Note 5.**

### Assessing false positive rate and statistical power

Genome-wide association analyses were conducted on the simulated data with 6 different methods. The simulated phenotypes were pre-adjusted by the top 10 PCs computed from a set of LD-pruned variants using flashPCA2^67^ (**Supplementary Note 6** and **Supplementary Figure 20)**. Since the number of variants was not large (556,269 variants after QC), we used all the variants to compute the sparse GRM for fastGWA and included all the variants as the “model SNPs” in the polygenic component for BOLT-LMM. After performing GWAS, we quantified the false positive rate, genomic inflation, and statistical power of each association method under each simulation setting.

### Real data analyses

We used fastGWA to perform a genome-wide association analysis for 2,173 traits in the UKB. We followed the QC pipeline as used in the Neale Lab’s UKB GWAS (**URLs**) with some modifications (e.g., we kept all the related individuals). In brief, only the participants with imputed genotype data and labelled as European ancestry (see the UKB Data-field 1001) were included in the association analyses (*n* = 456,422). Quantitative traits with >20% participants having the same phenotypic value and categorical traits were transformed into binary (TRUE/FALSE) or ordered-categorical variables, and the other quantitative traits were converted to z-scores by rank-based inverse normal transformation (INT). We removed binary traits with case fraction < 1% because the FPRs of MLM-based methods tend to be inflated for binary phenotypes with case fraction < 1% even for common variants^42^. For association analyses, we fitted age, age^2^, sex, age×sex, age^2^×sex, and the top 20 PCs provided by the UKB as covariates. The sex-specific traits were adjusted for age, age^2^, and the top 20 PCs. We excluded variants with MAF < 0.01 or missingness-rate > 0.1 for each trait. Clumping analysis was performed in a subset of 24 traits (listed in **Supplementary Table 3**) using PLINK2 with stringent criteria (bi-allelic variants only, *P*-value threshold = 5×10^−9^, window size = 5 Mb, and LD r^2^ threshold = 0.01), and a random subset of 10,000 unrelated individuals in the UKB was used as a LD reference panel.

We applied the same analysis pipeline used above to the WES data (152,327 variants with MAF ≥ 0.01 and 46,191 participants) for all the available traits in the UKB (excluding traits with *n* < 5000 or binary traits with case fraction < 1%, and variants with MAF < 0.01 or missingness-rate > 0.1 for each trait). We performed clumping analysis (bi-allelic variants only, *P*-value threshold = 0.05 / *m* with *m* being the number of variants tested for each trait, window size = 5 Mb, and LD r^2^ threshold = 0.01) of the fastGWA result for each of the 24 selected traits based on LD estimated from WES data of 42,974 unrelated individuals (estimated pairwise genetic relatedness < 0.05).

### URLs

fastGWA: http://cnsgenomics.com/software/gcta/#fastGWA

GCTA-GRM (computing GRM in biobank-scale data): http://cnsgenomics.com/software/gcta/#MakingaGRM

UKB GWAS summary statistics from fastGWA: http://cnsgenomics.com/software/gcta/#DataResource

Online tool to query the UKB summary statistics produced by fastGWA: http://fastgwa.info PheWeb: https://github.com/statgen/pheweb/

R-script to generate FAM: http://cnsgenomics.com/software/gcta/#fastGWA

UK Biobank: http://www.ukbiobank.ac.uk

PLINK2: https://www.cog-genomics.org/plink/2.0/

BOLT-LMM: https://data.broadinstitute.org/alkesgroup/BOLT-LMM/

LD score regression: https://github.com/bulik/ldsc

FlashPCA2: https://github.com/gabraham/flashpca

UKB Phenotype Processing scripts: https://github.com/Nealelab/UK_Biobank_GWAS

GeneATLAS: http://geneatlas.roslin.ed.ac.uk

UKB GWAS summary statistics from the Neale Lab: http://www.nealelab.is/uk-biobank

Eigen C++ library: http://eigen.tuxfamily.org/index.php?title=Main_Page

## Data availability

The individual-level genotype and phenotype data are available through formal application to the UK Biobank (**URLs**). All the summary-level statistics are available at our data portal (**URLs**).

## Acknowledgements

We thank Huanwei Wang and Julia Sidorenko for assistance in data preparation, Allan McRae for organising computing resources, Po-Ru Loh for constructive comments on the manuscript, the Neale Lab for making the data processing pipelines publicly available, and Alibaba Cloud - Australia and New Zealand for hosting the online tool. This research was supported by the Australian Research Council (DP160101343, DP160101056 and FT180100186), the Australian National Health and Medical Research Council (1078037, 1078901, 1113400 and 1107258), and the Sylvia & Charles Viertel Charitable Foundation. This study makes use of data from the UK Biobank (project ID: 12514). A full list of acknowledgments of this data set can be found in **Supplementary Note 8**.

## Author contributions

JY conceived and supervised the study. JY, LJ and ZZ designed the experiment. ZZ developed the software tools. LJ and ZZ performed the simulations and data analyses under the assistance and guidance from JY, PMV, TQ, NRW and KEK. PMV, NRW and JY contributed resources and funding. LJ and JY wrote the manuscript with the participation of all authors. All authors reviewed and approved the final manuscript.

