## Supplementary for "A resource-efficient tool for mixed model association analysis of large-scale data"

Jiang *et al.*

**Supplementary Note 1-8**

**Supplementary Tables 1-7**

**Supplementary Figures 1-20**

**Supplementary References**

### Supplementary Notes

#### Supplementary Note 1. Estimating the variance components by fastGWA-REML

Here we reiterate the fastGWA model

$$\boldsymbol{y} = \boldsymbol{x}_{snp}\beta_{snp} + \boldsymbol{X}_c\boldsymbol{\beta}_c + \boldsymbol{g} + \boldsymbol{e} \qquad \text{[S1]}$$

where $\boldsymbol{y}$ is an $n \times 1$ vector of mean centred phenotypes with $n$ being the sample size; $\boldsymbol{x}_{snp}$ is a vector of mean-centred genotype variables of a variant of interest with its effect $\beta_{snp}$; $\boldsymbol{X}_c$ is the incidence matrix of fixed covariates with their corresponding coefficients $\boldsymbol{\beta}_c$; $\boldsymbol{g}$ is a vector of the total genetic effects captured by pedigree relatedness with $\boldsymbol{g} \sim N(0, \boldsymbol{\pi}\sigma_g^2)$; $\boldsymbol{\pi}$ is the family relatedness matrix based on pedigree structure; $\boldsymbol{e}$ is a vector of residuals with $\boldsymbol{e} \sim N(0, \mathbf{I}\sigma_e^2)$. The variance-covariance matrix of $\boldsymbol{y}$ is $\boldsymbol{V} = \boldsymbol{\pi}\sigma_g^2 + \boldsymbol{I}\sigma_e^2$ and the generalized least squares estimate of $\beta_{snp}$ is $\hat{\beta}_{snp} = \frac{\boldsymbol{x}_{snp}^T\boldsymbol{V}^{-1}\boldsymbol{y}}{\boldsymbol{x}_{snp}^T\boldsymbol{V}^{-1}\boldsymbol{x}_{snp}}$ with $\text{var}(\hat{\beta}_{snp}) = \frac{1}{\boldsymbol{x}_{snp}^T\boldsymbol{V}^{-1}\boldsymbol{x}_{snp}}$. Therefore, to test whether $\beta_{snp} = 0$, we first need to estimate the variance components $\sigma_g^2$ and $\sigma_e^2$. As in most existing MLM-based association tools[1-7], to avoid running the variance estimation analysis repeatedly for each target variant, we estimate $\sigma_g^2$ and $\sigma_e^2$ under the null model

$$\boldsymbol{y} = \boldsymbol{X}_c\boldsymbol{\beta}_c + \boldsymbol{g} + \boldsymbol{e} \qquad \text{[S2]}$$

assuming the effect of a single variant on $\hat{\sigma}_g^2$ is negligible. The REML log-likelihood ($L$) function of model [2] can be written as

$$L = -\frac{1}{2}(\log|\boldsymbol{V}| + \log|\boldsymbol{X}_c{}^T\boldsymbol{V}^{-1}\boldsymbol{X}_c| + \boldsymbol{y}^T\boldsymbol{P}\boldsymbol{y}) \qquad \text{[S3]}$$

$$\text{with } \boldsymbol{P} = \boldsymbol{V}^{-1} - \boldsymbol{V}^{-1}\boldsymbol{X}_c(\boldsymbol{X}_c{}^T\boldsymbol{V}^{-1}\boldsymbol{X}_c)\boldsymbol{X}_c{}^T\boldsymbol{V}^{-1}$$

Conventional REML algorithms such as the average information (AI)[8] involve the computations of $\boldsymbol{V}^{-1}$, $\boldsymbol{P}$ and $\boldsymbol{P\pi}$, which is computationally intensive when $n$ is large even if $\boldsymbol{\pi}$ is sparse. Here we describe an algorithm (termed as fastGWA-REML) that uses grid search to estimate $\sigma_g^2$ without the need to compute $\boldsymbol{V}^{-1}$, $\boldsymbol{P}$ and $\boldsymbol{P\pi}$. For ease of computation, we first adjust the phenotype for covariates by linear regression (let $\boldsymbol{y}_{adj}$ denote a vector of phenotypes after adjustment). We can rewrite $L$ as $-\frac{1}{2}(\log|\boldsymbol{V}| + \log|\mathbf{1}^T\boldsymbol{V}^{-1}\mathbf{1}| + \boldsymbol{y}_{adj}^T\boldsymbol{V}^{-1}\boldsymbol{y}_{adj} - \boldsymbol{y}_{adj}^T\boldsymbol{V}^{-1}\mathbf{1}(\mathbf{1}^T\boldsymbol{V}^{-1}\mathbf{1})\mathbf{1}^T\boldsymbol{V}^{-1}\boldsymbol{y}_{adj})$ with $\mathbf{1}$ being an $n \times 1$ vector of 1's. All the elements in $L$ including $|\boldsymbol{V}|$, $\boldsymbol{V}^{-1}\mathbf{1}$ and $\boldsymbol{V}^{-1}\boldsymbol{y}_{adj}$ can be computed efficiently by the Cholesky decomposition of $\boldsymbol{V}$ (without the need of computing $\boldsymbol{V}^{-1}$) in sparse matrix setting. Because the computation of $L$ is extremely fast, we can use a grid search to obtain an estimate of $\sigma_g^2$ (note that $\hat{\sigma}_e^2$ can be computed as $\hat{\sigma}_P^2 - \hat{\sigma}_g^2$ with $\hat{\sigma}_P^2$ being the empirical variance of phenotype after adjustment).

The rationale underlying this grid-search method is similar to that in *Runcie et al.*[9]. We compute the log-likelihood scores given a grid of possible values of $\hat{\sigma}_g^2$ (e.g., $\hat{\sigma}_g^2 \in [0, 1.6\hat{\sigma}_P^2]$ with 100 steps, i.e., a step size of $0.016\hat{\sigma}_P^2$). Note that we define an upper limit to be large than $\hat{\sigma}_P^2$ to accommodate rare scenarios where the estimate of $\hat{\sigma}_g^2$ from the fastGWA model can be larger than $\hat{\sigma}_P^2$ if the true heritability is large in the presence of substantial common environmental effects. Next, we refine the search in a window around the $\hat{\sigma}_g^2$ value that produces the highest log-likelihood score (denoted by $\hat{\sigma}_{g(max)}^2$) with a window size of $0.2\hat{\sigma}_{g(max)}^2$ and 16 steps. For example, if $\hat{\sigma}_{g(max)}^2 = 0.16\hat{\sigma}_P^2$, we will refine the search in $\hat{\sigma}_g^2 \in [0.144\hat{\sigma}_P^2, 0.176\hat{\sigma}_P^2]$ with 16 steps (i.e., a step size of $0.002\hat{\sigma}_P^2$). We repeat this process iteratively until the difference in $\hat{\sigma}_g^2$ with the highest log-likelihood score between two adjacent iterations is smaller than $1 \times 10^{-10}\hat{\sigma}_P^2$. We confirmed by simulation that the estimates of $\sigma_g^2$ from fastGWA-REML were nearly identical to those from AI-REML implemented in GCTA [10] (the correlation between $\hat{\sigma}_g^2$ estimates from the two methods was > 0.9999; **Supplementary Figure 1**).

**Supplementary Note 2. Computing fastGWA test-statistics by the GRAMMAR-GAMMA approximation**

As mentioned in **Supplementary Note 1**, the effect of the target variant can be estimated by the generalised least squares approach as

$$\hat{\beta}_{snp} = \frac{\boldsymbol{x}_{snp}^T\boldsymbol{V}^{-1}\boldsymbol{y}}{\boldsymbol{x}_{snp}^T\boldsymbol{V}^{-1}\boldsymbol{x}_{snp}} \text{ with } \mathrm{var}(\hat{\beta}_{snp}) = \frac{1}{\boldsymbol{x}_{snp}^T\boldsymbol{V}^{-1}\boldsymbol{x}_{snp}} \qquad [S4]$$

In this equation, although $\boldsymbol{V}^{-1}\boldsymbol{x}_{snp}$ and $\boldsymbol{V}^{-1}\boldsymbol{y}$ can be computed efficiently using the Cholesky decomposition of $\boldsymbol{V}$ (precomputed in fastGWA-REML), $\boldsymbol{V}^{-1}\boldsymbol{x}_{snp}$ needs to be computed repeatedly for millions of variants across the genome, which is still a computational bottleneck. We can get around this problem by using the GRAMMAR-GAMMA approximation [11]. In brief, Equation [S4] can be rewritten as

$$\hat{\beta}_{snp} = \frac{\boldsymbol{x}_{snp}^T\boldsymbol{V}^{-1}\boldsymbol{y}}{\boldsymbol{x}_{snp}^T\boldsymbol{V}^{-1}\boldsymbol{x}_{snp}} = \left(\frac{\boldsymbol{x}_{snp}^T\boldsymbol{V}^{-1}\boldsymbol{y}}{\boldsymbol{x}_{snp}^T\boldsymbol{x}_{snp}}\right) / \left(\frac{\boldsymbol{x}_{snp}^T\boldsymbol{V}^{-1}\boldsymbol{x}_{snp}}{\boldsymbol{x}_{snp}^T\boldsymbol{x}_{snp}}\right) \qquad [S5]$$

In this equation, the first term $\frac{\boldsymbol{x}_{snp}^T\boldsymbol{V}^{-1}\boldsymbol{y}}{\boldsymbol{x}_{snp}^T\boldsymbol{x}_{snp}}$ is easy to compute as $\boldsymbol{V}^{-1}\boldsymbol{y}$ can be computed efficiently using the Cholesky decomposition, which only needs to be done once. The second term $\frac{\boldsymbol{x}_{snp}^T\boldsymbol{V}^{-1}\boldsymbol{x}_{snp}}{\boldsymbol{x}_{snp}^T\boldsymbol{x}_{snp}}$ appears to be dependent on $\boldsymbol{x}_{snp}$ in its form but results from previous studies[7,11] show that this term is nearly a constant (denoted by $\gamma$) regardless of SNP genotypes. In practice, we estimate $\gamma$ by the mean of $\frac{\boldsymbol{x}_{snp}^T\boldsymbol{V}^{-1}\boldsymbol{x}_{snp}}{\boldsymbol{x}_{snp}^T\boldsymbol{x}_{snp}}$ across 1,000 randomly selected variants that are not associated with the phenotype (excluding variants with linear regression $\chi^2 > 5$). As for the number of variants used to estimate $\gamma$, previous studies[7,12] suggest that 30 is sufficient. In our tool, since the computation of $\frac{\boldsymbol{x}_{snp}^T\boldsymbol{V}^{-1}\boldsymbol{x}_{snp}}{\boldsymbol{x}_{snp}^T\boldsymbol{x}_{snp}}$ is relatively fast, we use 1000 to ensure that $\hat{\gamma}$ is estimated with high precision (nevertheless, we have an option in our tool that allows users to change this number). Finally, we use $\hat{\gamma}$ to compute the summary statistics of all variants by this approximation

$$\hat{\beta}_{snp} \approx \frac{x_{snp}^T \boldsymbol{V}^{-1}\boldsymbol{y}}{x_{snp}^T x_{snp}} / \hat{\gamma} \text{ and } \mathrm{var}(\hat{\beta}_{snp}) \approx \frac{1}{x_{snp}^T x_{snp}} / \hat{\gamma} \qquad \text{[S6]}$$

We have confirmed by simulation that the test-statistics computed by this approximation were almost identical to those computed by the exact approach (**Supplementary Figure 2**). The computational complexity of equation [S6] is approximately $O(NM)$, nearly independent of the density of the sparse GRM. Note that the computational complexity of estimating $\hat{\gamma}$ (mainly attributed to the computation of $\boldsymbol{V}^{-1}\boldsymbol{x}_{snp}$) is approximately linearly proportional to sparse GRM density as indicated by the increase of runtime as a function of sparse GRM density in the fastGWA-REML analysis (**Supplementary Table 2**). Since $\hat{\gamma}$ only needs to be computed once under the null model, the effect of sparse GRM density on the overall runtime of the association test step is limited in a certain range.

**Supplementary Note 3. Constructing relatedness matrix from pedigree data and sparse GRM from SNP data**

If pedigree information is available, we can perform a fastGWA-Ped analysis with a relatedness matrix constructed from the expected relatedness coefficients (e.g., 0.5, 0.25, 0.125 and 0.0675 for the first-, second-, third- and fourth-degree relatives, respectively and 1 for monozygotic twins), similar to the traditional family-based MLMs[1,13-15]. We have provided an R-script (see **URLs**) to construct FAM based on unknown relatedness information (e.g., the inferred relatedness information provided by the UKB). Note that for fastGWA-Ped, to avoid singularity of matrix **V** for traits for which the estimate of residual variance component was negative or close to zero (e.g., < 10% of the phenotypic variance), the non-zero elements in the family relatedness matrix $\boldsymbol{\pi}$ can be recomputed from SNP data with minimal computing cost.

If the pedigree information is incomplete or is not available, the pedigree relatedness matrix can be replaced by a sparse GRM. However, computing a GRM is time-consuming with a time complexity of $O(MN^2)$ when the sample size is large. We have developed a new GRM function in GCTA using the BLAS packages available in the Intel MKL library to compute a GRM in a very efficient manner via bitwise operations (**URLs**). We have also provided a shell script in the GCTA website (**URLs**) to divide the whole computation process into a number of computing jobs to be parallelized in a high-performance computing system.

In the fastGWA analysis of the UKB data, we computed the GRM for 456,422 individuals of European ancestry using 565,631 slightly LD-pruned HapMap3[16] variants (LD-pruning parameters used in PLINK: window size = 1000kb, step size = 100, $r^2$ = 0.9 and MAF ≥ 0.01). Note that this set variants are sufficient to capture the relatedness among close relatives (e.g., those with genetic relatedness > 0.05). In the fastGWA-ped analysis, we inferred the pedigree relationships based on the KING[17,18] relatedness estimates provided by the UKB. The runtime to build a GRM for the full UKB cohort was around 2.2 hours with 100 jobs (each job was assigned with 8 GB memory and 6 CPUs). There is an efficient tool in GCTA to convert a full-dense GRM to a sparse GRM given a relatedness threshold (default value = 0.05 but can be changed by users) and the computing cost of this conversion is low even for large data sets like the UKB. There is also an option in GCTA that allows users to output a sparse GRM on-the-fly without storing the full dense GRM. Note that the same set of variants were used in the BOLT-REML and GCTA-GREML analyses to estimate the "genetic variance" and/or variance due to common environmental effect (see **Discussion** and **Figure 3**).

We noticed that there were some discrepancies between the pedigree relatedness inferred from relatedness estimates provided by the UKB and our sparse GRM (**Supplementary Figure 18**). This is mainly because we used GRM to pick up more distant related pairs with relatedness coefficients between 0.05 to 0.125. In contrast, the relatedness estimates from the UKB were evaluated according to a more sophisticated strategy (e.g., using a different set of markers with $m$ = 93,511 and excluding a small proportion of individuals with higher missingness rates)[18]. The number of related pairs provided by the UKB was 107,162 (no further than third-degree relatives) involving 147,731 unique individuals, while the number of related pairs based on our sparse GRM (estimated from 565,631 common HapMap3 variants) was 175,708 with 212,018 unique individuals. These two relatedness estimates were used for two different primary purposes: the UKB estimate was primarily used to make explicit inference about the family relatedness among samples, while our estimate was used to capture the relatedness between all close and distant relatives with the primary aim to control for the confounding in association test.

**Supplementary Note 4. Genotype data used in simulations**

To generate a cohort with relatedness and substantial population stratification, we sampled segments of SNP genotypes from existing GWAS data based on a mosaic simulation scheme modified from Loh *et al.*[7]. Detailed procedures have been listed below (see **Supplementary Figure 5** for a schematic diagram):

1) Randomly selecting two groups of individuals with different ancestry backgrounds from the UKB participants. Based on the self-reported ethnic background, we first extracted all the

UKB participants reported as “British” and “Irish” (see Data Field 21000 of UKB at http://biobank.ctsu.ox.ac.uk/crystal/field.cgi?id=21000). However, the self-reported ancestry may not be accurate, as we observed inconsistency between one’s belief and his/her actual genetic background estimated from genotype data (**Supplementary Figure 3)**. Thus, we selected 9,000 unrelated self-reported “British” and 9,000 unrelated “Irish” participants with relatively large differences in the first two PCs (**Supplementary Figure 4**) to ensure sufficient genetic separation between the two groups. These individuals were the “ancestors” (or founders) of our simulated individuals in step 2.

2) Generating genotypes of 90,000 unrelated individuals (i.e., 45,000 unrelated “British” and 45,000 unrelated “Irish”). To generate 45,000 unrelated “British” individuals, we first divided the genomes (536,684 variants in total) of all 9,000 “British” ancestors into 269 consecutive segments of approximately 2,000 variants. Then, to simulate the genotype of one individual, 100 ancestors were randomly sampled from the 9,000 “British” ancestors. Next, we randomly selected each segment from one of the 100 ancestors, and aligned all the 269 sampled segments back together to form a new complete genome. This would be one simulated “British” individual. By repeating these steps, 45,000 unrelated “British” individuals were generated. We used the same strategy to generate 45,000 unrelated “Irish” individuals by sampling the segments from the “Irish” ancestors.

3) Generating genotypes of 10,000 related individuals (i.e., 5,000 related “British” and 5,000 related “Irish”). A similar scheme as in step 2) was applied to generate related individuals. We define “related individuals” as those who are related with at least one other individual in the sample with genetic relatedness ≥ 0.05. Two individuals, as one related pair, were generated simultaneously each time. To mimic different degrees of relatedness, the segments for each pair of 1st degree of relatives were randomly sampled from 2 “common ancestors”, and the segments for each pair of 2nd degree of relatives were randomly sampled from 4 “common ancestors”. Instead of using the original 18,000 ancestors from step 1), we generated an independent set of additional 10,000 “British” individuals and 10,000 “Irish” individuals as the ancestors for these related individuals. A total of 2,500 pairs of 1st degree of relatives (genetic relatedness = ~0.5; 1,250 “British” pairs and 1,250 “Irish” pairs) and 2,500 pairs of 2nd degree of relatives (genetic relatedness = ~0.25; 1,250 “British” pairs and 1,250 “Irish” pairs) were eventually simulated.

In summary, we generated genotype data of two groups of individuals with reasonably large difference in genetic ancestry between the two groups and substantial proportion of closely related individuals within each group. The difference in ancestry allowed us to simulate the effect of population stratification and the related individuals allowed us to simulate shared environmental effects.

We also performed simulations based on real genotypes from the UKB (see **Methods** section in the main text and **Supplementary Note 5** for the method to generate phenotypes). We extracted SNP array genotyped data of 565,631 variants on 100,000 individuals of European ancestry. The individuals were not selected at random but purposely sampled to achieve a higher sparse GRM density (85,727 unique pairs of individuals with estimated genetic relatedness > 0.05; sparse GRM density = $2.7 \times 10^{-5}$) than that in the whole UKB sample (sparse GRM density =$3.9\times 10^{-6}$).

**Supplementary Note 5. Strategies for simulating phenotypes**

We used a set of different parameters to simulate phenotypes based on the simulated/real genotypes by the following model:

$$\boldsymbol{y} = \boldsymbol{g} + \boldsymbol{z}b_p + \boldsymbol{e}_c + \boldsymbol{e} \quad [S7]$$

1) $\boldsymbol{g} = \sum_{i=1}^{m} \boldsymbol{x}_i b_i$ where $\boldsymbol{x_i}$ is a vector of genotypes of the $i^{th}$ causal variant across all individuals with its effect $b_i$ generated from a standard normal distribution $N(0, 1)$. The number of causal variants ($m$) was set to 10,000 to mimic a polygenic trait, all of which were randomly sampled from variants on the <u>odd</u> chromosomes (leaving the variants on the <u>even</u> chromosomes to quantify the inflation in test-statistics under the null). We also tested the methods with more causal variants, i.e., 20,000, 40,000, or 80,000.
2) $\boldsymbol{z}$ is an indicator vector consists of 0 (indicating "British") and 1 (indicating "Irish") with $b_p$ being the mean difference in phenotype between the two groups. The value of $b_p$ does not matter as $\boldsymbol{z}b_p$ is standardised in the final step, and any positive value of $b$ yields the same result. The purpose of this step is to simulate population stratification effect by creating a phenotypic mean difference between the two ancestry groups.
3) $\boldsymbol{e}_c$ is a vector of shared environmental effects generated by a) identifying "close relatives" (including the simulated first- and second-degree relatives as well as those pairs with estimated genetic relatedness > 0.05; **Supplementary Note 4**) and grouping them into "families" and extended "families"; b) randomly sampling an effect from a normal distribution to all the individuals in each family.
4) $\boldsymbol{e}$ is a vector of residual effects, randomly generated from a normal distribution $\boldsymbol{e}{\sim}N(0,1)$.

The phenotypic value for each sample was a weighted sum of all the standardised components above. The phenotypic variance ($V_p$) was set to 1. The weights were the square root of the variance proportion of each component, determined by the following: a) genetic variance: $V_g = 0.4 \times V_p$; b) variance due to population stratification: $V_{pop} = 0.05 \times V_p$; c) variance due to common environmental effects: $V_{related} = 0.1\ or\ 0.2\ \times V_p$ (for related individuals); and d) residual variance: $V_{residual} = V_p - V_g - V_{pop} - V_{related}$.

Additional simulations were conducted for phenotypes that do not follow a normal distribution including binary phenotypes, exponentially transformed phenotypes, and phenotypes with residuals following a $\chi_1^2$ distribution.

1) For binary phenotypes, we simulated quantitative phenotypes for the 100,000 simulated individuals as described in **Supplementary 4**, and dichotomized the phenotypes given three sample prevalence rates (i.e., 0.3, 0.1, or 0.01).
2) For exponentially transformed phenotypes, we simulated the original quantitative phenotypes ($y'$) as described above and transformed them using a natural exponential function $y = e^{y'}$.
3) For phenotypes with residuals following a $\chi_1^2$ distribution, we simulated the genetic effect ($\boldsymbol{g}$), the effect of population stratification ($\boldsymbol{z}\boldsymbol{b}_p$), and the common environmental effect among relatives ($\boldsymbol{e}_c$) using the same strategies as described in **Supplementary 4**. We then generated the residual effect ($\boldsymbol{e}$) from a chi-squared distribution with 1 degree of freedom (i.e., $\boldsymbol{e} \sim \chi_1^2$).

For simulations based on real genotypes described in **Supplementary Note 5** and **Supplementary Figure 13**, we used the same strategies to generate the phenotypes except that we did not simulate the effect of population stratification ($\boldsymbol{z}\boldsymbol{b}_p$). This is because the ancestry background in real genotype data was more complicated than that in the simulated genotype data and the self-reported ancestry information was not reliable enough for simulation (**Supplementary Figure 3**).

**Supplementary Note 6. Principal component analysis**

We compared three different principal component analysis (PCA) methods using our simulated genotype data, namely flashPCA2 (or pruned PCA, with a recommended pruning step and a projection step, see **URL** and Ref.[19]), exact PCA (implemented in GCTA using all the variants without pruning, see Ref.[20]), and projection PCA (proj. PCA, implemented in GCTA). The proj. PCA method can be described as: 1) randomly extracting 10,000 individuals from the sample ($n$ = 100,000); 2) performing exact PCA using the subset of individuals and estimating the loadings of each SNP to the top PCs; and 3) computing the top PCs of the whole sample based on the SNP loadings estimated from the subset.

The first and second PCs calculated from flashPCA2, as well as those from exact PCA, were plotted in **Supplementary Figure 6**. Apart from the scale difference, both methods managed to separate the individuals into two groups based on the first two PCs. We examined the proportion of phenotypic variance ($V_p$) explained by the top PCs. There was no significant difference among all three methods (**Supplementary Figure 20**), as all of them accounted for approximately 5% of $V_p$, consistent with the parameter used to simulate data. However, fitting PCs calculated from all the variants (by exact PCA) would lead to slight deflation of test statistics (**Supplementary Figure 19**). This is because each PC is essentially a feature extracted from the variants, and fitting PCs as covariates while testing a target SNP is equivalent to fitting the target SNP more than once in the model, leading to deflated test-statistics under the null. If PCs are computed using a set of LD-pruned variants with a relatively stringent threshold, the target SNP is much less likely to be included in computing the PCs. In this case, the test statistics are less likely to be deflated under the null (**Figure 1a)**. We therefore adjusted the phenotypes by the top 10 PCs from flashPCA2 (76,103 LD-pruned variants with window size = 1 Mb, step size = 50 variants, and LD $r^2$ threshold = 0.05 as recommended by flashPCA2) in all the subsequent association tests in the simulation study. The number of PCs used is justified by the result that the top 10 PCs were sufficient to capture the majority of phenotype variation due to population stratification (**Supplementary Figure 20**).

In real data analysis, we adjusted each of the 2,173 UKB traits by the top 20 PCs provided by the UKB data analysis team[18] (computed by fastPCA[21] using a similar strategy as in flashPCA2).

**Supplementary Note 7. LD scores**

LD score of a SNP is defined as the sum of LD $r^2$ between the target SNP and all the other variants in a genomic region adjusting for chance correlations[22]. LD scores are required for both BOLT-LMM (mixture model) and LD score regression (LDSC) analyses. In real data analysis, we used LD scores provided by BOLT-LMM and LDSC software tools (computed from SNP data of individuals of European ancestry in the 1000 Genomes Project[23]; see **URLs**). For simulations based on the simulated genotype (**Supplementary Note 4**), we computed LD scores from the unrelated individuals in the sample (excluding one of each pair of individuals with estimated relatedness < 0.05) using GCTA[24] (with window size of 1 Mb, 10 Mb or 20 Mb). For simulations based on real genotype, two sets of LD scores were used for comparison (**Supplementary Figure 17b**): 1) LD scores provided at the BOLT-LMM website (https://data.broadinstitute.org/alkesgroup/BOLT-LMM/), and 2) LD scores computed from the unrelated individuals in the sample.

**Supplementary Note 8. Acknowledgements**

**UKB:** This study has been conducted using UK Biobank resource under Application Number 12514. UK Biobank was established by the Wellcome Trust medical charity, Medical Research Council, Department of Health, Scottish Government and the Northwest Regional Development Agency. It has also had funding from the Welsh Assembly Government, British Heart Foundation and Diabetes UK.

### Supplementary Tables

**Supplementary Table 1.** Comparison of runtime between GCTA-fastGWA, BOLT-LMM, and PLINK2 tested on hard disk drives (HDD). The data used in this test consisted of 8,531,416 common variants, of which 565,631 LD-pruned variants were used as "model SNPs" in BOLT-LMM (**Supplementary Note 3**). The runtime of fastGWA or BOLT-LMM consists of two steps: a) the estimation of mixed model parameters ("Para. Est."), and b) the association test ("Assoc."). The PLINK2 linear regression was performed using all the individuals. All tests were performed the same computing environment: 96 GB memory and 16 CPU cores with hard disk drives in one computer node.

| Sample Size | GCTA-fastGWA v1.92.3 | | | BOLT-LMM v2.3.2 | | | PLINK2 v2.00a2 |
|---|---|---|---|---|---|---|---|
| | Para. Est. (h) | Assoc. (h) | Total (h) | Para. Est. (h) | Assoc. (h) | Total (h) | Total (h) |
| 50,000 | 0.05 | 0.15 | 0.20 | 0.94 | 1.17 | 2.11 | 0.24 |
| 100,000 | 0.04 | 0.24 | 0.28 | 2.21 | 2.28 | 4.49 | 0.54 |
| 200,000 | 0.02 | 0.39 | 0.41 | 5.72 | 4.62 | 10.34 | 0.97 |
| 300,000 | 0.02 | 0.48 | 0.50 | 9.73 | 6.71 | 16.44 | 1.39 |
| 400,000 | 0.02 | 0.67 | 0.69 | 15.59 | 9.27 | 24.86 | 2.12 |

**Supplementary Table 2**. Computational costs of fastGWA-REML and fastGWA-HE regression with different levels of sparse GRM density. The total sample size was fixed to be 400,000. Sparse GRM density is defined as the proportion of non-zero elements in a sparse GRM. The computing environment is the same as that in analysis presented in Table 1: 96 GB memory and 16 CPU cores with solid-state disk in one computer node. The last row shows the actual sparse GRM density and runtime of the whole UKB data ($n$ = 456,422). "sec" = second and "GB" = gigabyte.

| Density | Number of unique related pairs | REML runtime (sec) | REML Memory usage (GB) | REML Virtual Memory usage (GB) | HE-Reg runtime (sec) | HE-Reg Memory usage (GB) | HE-Reg Virtual Memory usage (GB) |
|---|---|---|---|---|---|---|---|
| $5.0 \times 10^{-6}$ | 200,000 | 12.2 | 2.95 | 4.72 | 12.7 | 1.88 | 3.85 |
| $7.5 \times 10^{-6}$ | 400,000 | 17.5 | 3.33 | 5.67 | 12.6 | 1.88 | 4.00 |
| $1.0 \times 10^{-5}$ | 600,000 | 22.3 | 4.03 | 6.71 | 12.3 | 1.89 | 3.88 |
| $1.25 \times 10^{-5}$ | 800,000 | 26.4 | 4.65 | 7.69 | 12.7 | 1.90 | 3.89 |
| $1.5 \times 10^{-5}$ | 1,000,000 | 28.8 | 4.61 | 7.62 | 12.3 | 1.87 | 3.90 |
| $2.5 \times 10^{-5}$ | 1,800,000 | 45.6 | 5.51 | 7.53 | 12.3 | 1.94 | 3.92 |
| $5.0 \times 10^{-5}$ | 3,800,000 | 63.1 | 9.98 | 12.96 | 12.4 | 2.01 | 4.03 |
| $1.0 \times 10^{-4}$ | 7,800,000 | 140.1 | 16.51 | 22.12 | 12.5 | 2.24 | 4.44 |
| $3.9 \times 10^{-6}$ (UKB) | 175,708 | 8.8 | 2.95 | 4.99 | 12.6 | 1.89 | 3.85 |

**Supplementary Table 3.** Abbreviated names of the 24 UKB quantitative traits. The first column is the abbreviation, the second column is the data-field ID of each trait, the third column is the number of records available for analysis, the fourth column is the type of phenotype data (integer or continuous variable), the fifth column contains the full names of these phenotypes, and the last column indicates which traits are female-specific.

| Trait Abbr. | Data-field | Count | Type | Description | Note |
|---|---|---|---|---|---|
| WC | 48 | 500,376 | Continuous | Waist circumference | |
| HC | 49 | 500,317 | Continuous | Hip circumference | |
| HT | 50 | 499,997 | Continuous | Standing height | |
| WT | 21002 | 499,762 | Continuous | Weight | |
| BMI | 21001 | 499,431 | Continuous | Body mass index | |
| HGSR | 47 | 499,193 | Integer | Hand grip strength (right) | |
| HGSL | 46 | 499,126 | Integer | Hand grip strength (left) | |
| MTCIM | 20023 | 496,709 | Integer | Mean time to correctly identify matches | |
| BMR | 23105 | 492,388 | Continuous | Basal metabolic rate | |
| BFP | 23099 | 492,127 | Continuous | Body fat percentage | |
| DBP | 4079 | 472,416 | Integer | Diastolic blood pressure, automated reading | |
| SBP | 4080 | 472,411 | Integer | Systolic blood pressure, automated reading | |
| FVC | 3062 | 453,724 | Continuous | Forced vital capacity | |
| FEV | 3063 | 453,724 | Continuous | Forced expiratory volume in 1-second | |
| PEF | 3064 | 453,724 | Integer | Peak expiratory flow | |
| NTS | 20127 | 401,596 | Integer | Neuroticism score | |
| EA | 845 | 336,769 | Integer | Age completed full time education | |
| hBMD | 78 | 279,104 | Continuous | Heel bone mineral density (BMD) T-score, automated | |
| BW | 20022 | 277,009 | Continuous | Birth weight | |
| AMena | 2714 | 272,927 | Integer | Age at menarche | Female-specific factors |
| AFLB | 2754 | 184,987 | Integer | Age at first live birth | Female-specific factors |
| PR | 4194 | 170,759 | Integer | Pulse rate | |
| FIS | 20016 | 165,471 | Integer | Fluid intelligence score | |
| AMeno | 3581 | 165,363 | Integer | Age at menopause (last menstrual period) | Female-specific factors |

**Supplementary Table 4.** LD score regression attenuation ratio (SE) for the 24 UKB traits. Phenotypes are ordered by descending sample size (*n*). The GWAS summary statistics are from the fastGWA and BOLT-LMM-Inf analyses in this study, the Neale Lab, and the GeneATLAS, respectively. "\" represents that the trait is not available in GeneATLAS. The abbreviated and full names of the traits are listed in **Supplementary Table 1**.

| Trait | Attenu. ratio (fastGWA) | *n* (fastGWA) | Attenu. ratio (Neale Lab) | *n* (Neale Lab) | Attenu. ratio (GeneATLAS) | Attenu. ratio (BOLT-LMM-Inf) |
|---|---|---|---|---|---|---|
| WC | 0.0706 (0.0087) | 455,545 | 0.0712 (0.0104) | 360,564 | 0.0707 (0.008) | 0.0775 (0.0081) |
| HC | 0.0854 (0.0085) | 455,495 | 0.0865 (0.0098) | 360,521 | 0.0867 (0.0079) | 0.0959 (0.008) |
| HT | 0.1164 (0.0081) | 455,332 | 0.1206 (0.0089) | 360,388 | 0.112 (0.0068) | 0.1195 (0.0069) |
| WT | 0.0841 (0.0079) | 455,010 | 0.0838 (0.0091) | 360,116 | 0.0845 (0.0075) | 0.0923 (0.0075) |
| BMI | 0.0695 (0.0075) | 454,841 | 0.0715 (0.0086) | 359,983 | 0.0683 (0.0072) | 0.0782 (0.0071) |
| HGSR | 0.0746 (0.0108) | 454,473 | 0.0777 (0.0127) | 359,729 | 0.0658 (0.0096) | 0.0783 (0.0104) |
| HGSL | 0.0681 (0.0111) | 454,417 | 0.0672 (0.0125) | 359,704 | 0.0602 (0.0102) | 0.0695 (0.0108) |
| MTCIM | 0.0533 (0.0138) | 453,043 | 0.052 (0.0163) | 358,695 | 0.0472 (0.0154) | 0.0572 (0.0138) |
| BMR | 0.0983 (0.0083) | 448,348 | 0.1001 (0.0093) | 354,825 | 0.0971 (0.0079) | 0.1047 (0.0076) |
| BFP | 0.0808 (0.0081) | 448,114 | 0.0791 (0.0093) | 354,628 | 0.0836 (0.0076) | 0.0911 (0.0079) |
| DBP | 0.0892 (0.0114) | 430,029 | 0.0836 (0.0129) | 340,162 | \ | 0.0906 (0.0111) |
| SBP | 0.0901 (0.0102) | 430,025 | 0.0828 (0.012) | 340,159 | \ | 0.0923 (0.0097) |
| FVC | 0.1121 (0.0096) | 415,931 | 0.1055 (0.0104) | 329,404 | \ | 0.1163 (0.0097) |
| FEV | 0.0976 (0.0096) | 415,931 | 0.0891 (0.0101) | 329,404 | \ | 0.1033 (0.0094) |
| PEF | 0.0874 (0.0134) | 415,931 | 0.0852 (0.0155) | 329,404 | \ | 0.0928 (0.0131) |
| NTS | 0.0441 (0.0129) | 369,407 | 0.0443 (0.0155) | 293,006 | \ | 0.0501 (0.0125) |
| EA | 0.0918 (0.0156) | 304,998 | 0.084 (0.0171) | 240,547 | \ | 0.1043 (0.0153) |
| hBMD | 0.1139 (0.011) | 262,294 | 0.1147 (0.0119) | 206,589 | \ | 0.1184 (0.0105) |
| BW | 0.1087 (0.0228) | 258,857 | 0.1113 (0.0244) | 205,475 | \ | 0.1307 (0.0221) |
| AMena | 0.0337 (0.0108) | 240,378 | 0.038 (0.0119) | 188,644 | \ | 0.0402 (0.0104) |
| AFLB | 0.07 (0.0172) | 168,097 | 0.0733 (0.0186) | 131,987 | \ | 0.0743 (0.018) |
| PR | 0.0737 (0.0237) | 149,082 | 0.0687 (0.0284) | 118,850 | \ | 0.0837 (0.0229) |
| FIS | 0.053 (0.015) | 146,808 | 0.058 (0.018) | 117,131 | \ | 0.0496 (0.0138) |
| AMeno | 0.0365 (0.0315) | 141,926 | 0.0321 (0.0366) | 111,593 | \ | 0.0497 (0.0314) |
| **Mean (the first 10 traits)** | 0.0801 | | 0.0810 | | 0.0776 | 0.0864 |
| **Mean (all)** | 0.0792 | | 0.0783 | | \ | 0.0859 |

**Supplementary Table 5.** Number of near-independent genome-wide significant associations from GWAS analyses of the imputed data for 24 quantitative traits in the UKB. The near-independent signals were from PLINK clumping analysis (MAF ≥ 0.01, *P*-value threshold = $5\times10^{-9}$, window size = 5 Mb, and LD $r^2$ threshold = 0.01) of the GWAS summary statistics produced by four association methods, fastGWA, LR-unRel (results from the Neale Lab), DISSECT-LOCO (results from GeneATLAS), and BOLT-LMM-Inf. The sample size used in the BOLT-LMM-Inf analysis was the same as that in the fastGWA analysis. Phenotypes are ordered by descending sample size (*n*). The abbreviated and full names of the traits can be found in **Supplementary Table 1**.

| Trait | No. signals (fastGWA) | *n* (fastGWA) | No. signals (Neale Lab) | *n* (Neale Lab) | No. signals (GeneATLAS) | No. signals (BOLT-LMM-Inf) |
|---|---|---|---|---|---|---|
| WC | 399 | 455,545 | 246 | 360,564 | 433 | 449 |
| HC | 468 | 455,495 | 331 | 360,521 | 516 | 572 |
| HT | 1,611 | 455,332 | 1,318 | 360,388 | 2,259 | 2,268 |
| WT | 620 | 455,010 | 449 | 360,116 | 736 | 809 |
| BMI | 534 | 454,841 | 368 | 359,983 | 594 | 681 |
| HGSR | 161 | 454,473 | 94 | 359,729 | 197 | 172 |
| HGSL | 142 | 454,417 | 84 | 359,704 | 170 | 148 |
| MTCIM | 52 | 453,043 | 36 | 358,695 | 38 | 57 |
| BMR | 807 | 448,348 | 602 | 354,825 | 914 | 1,051 |
| BFP | 451 | 448,114 | 305 | 354,628 | 514 | 545 |
| DBP | 253 | 430,029 | 174 | 340,162 | \ | 280 |
| SBP | 282 | 430,025 | 211 | 340,159 | \ | 331 |
| FVC | 388 | 415,931 | 271 | 329,404 | \ | 436 |
| FEV | 458 | 415,931 | 329 | 329,404 | \ | 524 |
| PEF | 137 | 415,931 | 90 | 329,404 | \ | 145 |
| NTS | 64 | 369,407 | 46 | 293,006 | \ | 75 |
| EA | 37 | 304,998 | 20 | 240,547 | \ | 38 |
| hBMD | 460 | 262,294 | 358 | 206,589 | \ | 567 |
| BW | 119 | 258,857 | 75 | 205,475 | \ | 124 |
| AMena | 182 | 240,378 | 124 | 188,644 | \ | 203 |
| AFLB | 20 | 168,097 | 10 | 131,987 | \ | 17 |
| PR | 70 | 149,082 | 50 | 118,850 | \ | 71 |
| FIS | 39 | 146,808 | 23 | 117,131 | \ | 51 |
| AMeno | 85 | 141,926 | 62 | 111,593 | \ | 90 |
| **Sum (the first 10 traits)** | **5,245** | | **3,833** | | **6,371** | **6,752** |
| **Sum (all traits)** | **7,839** | | **5,676** | | **** | **9,704** |

**Spplementary Table 6.** Number of exome-wide significant associations from the fastGWA analysis of the WES data for 24 quantitative traits in the UKB. Phenotypes are ordered by descending sample size (*n*). The abbreviated and full names of the traits can be found in **Supplementary Table 1**. Clumping analysis criteria: *P*-value threshold = 0.05/ the number of tested variants, window size = 5 Mb, and LD $r^2$ threshold = 0.01. Conditional fastGWA: fastGWA analysis of a WES variant conditioning on the GWAS signals (with 10 Mb of the WES variant) identified from the imputed data.

| **Trait** | ***n*** | **fastGWA** | **Conditional fastGWA** |
|---|---|---|---|
| WC | 46135 | 0 | \ |
| HC | 46133 | 5 | 0 |
| HT | 46116 | 64 | 0 |
| WT | 46067 | 12 | 0 |
| BMI | 46051 | 8 | 0 |
| MTCIM | 46025 | 1 | 1 |
| HGSL | 45973 | 0 | \ |
| HGSR | 45949 | 1 | 0 |
| DBP | 45660 | 2 | 0 |
| SBP | 45659 | 4 | 0 |
| BMR | 45172 | 20 | 1 |
| BFP | 45143 | 3 | 0 |
| FEV | 41430 | 3 | 0 |
| FVC | 41430 | 5 | 0 |
| PEF | 41430 | 1 | 0 |
| NTS | 38071 | 0 | \ |
| FIS | 37351 | 6 | 1 |
| PR | 37166 | 5 | 1 |
| EA | 29619 | 0 | \ |
| BW | 27450 | 0 | \ |
| AMena | 24450 | 0 | \ |
| AFLB | 16703 | 0 | \ |
| AMeno | 14088 | 7 | 0 |
| hBMD | 7066 | 1 | 0 |
| **Total** | **** | **148** | **4** |

**Supplementary Table 7.** Summary statistics of the exome-wide significant associations from the fastGWAS analysis of the UKB WES data (*n* = 46,191) for 24 quantitative traits. PLINK clumping criteria: *P*-value threshold = 0.05/the number of variants tested for a trait, window size = 5 Mb, and LD $r^2$ threshold = 0.01. Each variant is named in a format "Chromosome:Position:Allele 1:Allele 2" based on the Genome Reference Consortium Human Build 38. "EA" = effect allele; "Freq." = frequency of the effect allele; *n* = sample size; "beta" = estimated variant effect; "se" = standard error of the estimated variant effect; p = p-value. Shown are also the WES GWAS summary statistics from the fastGWA analysis conditioning on GWAS signals (within 10Mb of the WES variant in either direction) identified from the analysis of the whole UKB imputed data (*n* = 456,422).

| Trait | Variant | EA | Freq. | *n* | fastGWA | | | Conditional fastGWA | | |
|---|---|---|---|---|---|---|---|---|---|---|
| | | | | | beta | se | p | beta | se | p |
| AMeno | 1:38895943:G:A | A | 0.457 | 14083 | -0.064 | 0.0113 | 1.44E-08 | -0.028 | 0.0114 | 0.015 |
| AMeno | 19:55319820:A:G | G | 0.391 | 14088 | -0.084 | 0.0118 | 8.53E-13 | -0.009 | 0.0119 | 0.441 |
| AMeno | 4:83446133:G:A | A | 0.476 | 14075 | 0.061 | 0.0111 | 4.61E-08 | 0.004 | 0.0111 | 0.710 |
| AMeno | 6:10887043:C:G | C | 0.174 | 14088 | 0.078 | 0.0150 | 1.80E-07 | 0.002 | 0.0156 | 0.911 |
| AMeno | 12:66310445:A:G | G | 0.031 | 14088 | 0.214 | 0.0331 | 8.82E-11 | 0.002 | 0.0333 | 0.960 |
| AMeno | 8:38030499:C:G | C | 0.225 | 14087 | 0.084 | 0.0136 | 6.66E-10 | 0.000 | 0.0136 | 0.974 |
| AMeno | 20:5967581:G:A | A | 0.061 | 14088 | 0.232 | 0.0238 | 1.35E-22 | 0.000 | 0.0238 | 1.000 |
| BFP | 20:63738996:T:C | T | 0.340 | 45131 | -0.027 | 0.0051 | 9.07E-08 | -0.003 | 0.0051 | 0.574 |
| BFP | 2:24918669:A:G | G | 0.484 | 45143 | 0.027 | 0.0050 | 5.69E-08 | -0.001 | 0.0051 | 0.872 |
| BFP | 11:27700751:G:T | T | 0.319 | 45143 | 0.028 | 0.0054 | 2.87E-07 | -0.001 | 0.0054 | 0.911 |
| BMI | 11:27700751:G:T | T | 0.319 | 46051 | 0.038 | 0.0069 | 5.30E-08 | 0.024 | 0.0070 | 0.0007 |
| BMI | 2:24918669:A:G | G | 0.484 | 46051 | 0.039 | 0.0065 | 2.53E-09 | -0.001 | 0.0065 | 0.891 |
| BMI | 15:67824962:T:A | A | 0.225 | 46051 | -0.041 | 0.0077 | 9.30E-08 | 0.001 | 0.0080 | 0.914 |
| BMI | 3:49860567:A:G | A | 0.490 | 46051 | -0.033 | 0.0065 | 2.57E-07 | 0.001 | 0.0067 | 0.938 |
| BMI | 18:60372043:C:T | T | 0.021 | 46051 | -0.121 | 0.0229 | 1.29E-07 | -0.001 | 0.0232 | 0.973 |
| BMI | 1:177929986:G:C | C | 0.206 | 46051 | 0.047 | 0.0080 | 4.98E-09 | 0.000 | 0.0081 | 0.989 |
| BMI | 19:45678134:G:C | C | 0.194 | 46051 | -0.044 | 0.0082 | 6.23E-08 | 0.000 | 0.0083 | 0.994 |
| BMI | 4:25407216:G:A | A | 0.233 | 46051 | -0.046 | 0.0077 | 1.56E-09 | 0.000 | 0.0078 | 0.996 |
| BMR | 12:6678624:A:C | C | 0.056 | 45172 | 0.051 | 0.0095 | 9.18E-08 | 0.059 | 0.0102 | 4.93E-09 |
| BMR | 3:129268887:G:A | A | 0.101 | 45165 | 0.037 | 0.0070 | 9.60E-08 | 0.034 | 0.0071 | 2.0E-6 |
| BMR | 2:23703363:C:T | T | 0.130 | 45172 | -0.037 | 0.0063 | 7.30E-09 | -0.015 | 0.0064 | 0.021 |
| BMR | 2:36581207:C:T | T | 0.351 | 45167 | 0.022 | 0.0044 | 2.90E-07 | 0.007 | 0.0045 | 0.143 |
| BMR | 4:17845658:A:C | C | 0.119 | 45063 | -0.037 | 0.0065 | 1.10E-08 | -0.008 | 0.0066 | 0.210 |
| BMR | 7:92618019:A:G | G | 0.250 | 45172 | 0.027 | 0.0049 | 3.43E-08 | 0.005 | 0.0050 | 0.288 |
| BMR | 9:108897159:T:C | C | 0.049 | 45172 | 0.056 | 0.0098 | 1.15E-08 | 0.009 | 0.0105 | 0.380 |
| BMR | 6:130060101:T:C | T | 0.311 | 45172 | 0.030 | 0.0046 | 1.20E-10 | 0.002 | 0.0047 | 0.681 |
| BMR | 1:155041950:G:A | A | 0.202 | 45172 | 0.028 | 0.0053 | 7.05E-08 | 0.002 | 0.0054 | 0.715 |
| BMR | 16:2105296:A:G | G | 0.169 | 45172 | -0.030 | 0.0057 | 1.46E-07 | -0.002 | 0.0059 | 0.749 |
| BMR | 4:145159425:D:1 | A | 0.359 | 45171 | 0.024 | 0.0044 | 3.33E-08 | 0.001 | 0.0045 | 0.771 |
| BMR | 17:63930138:A:G | A | 0.354 | 45172 | -0.025 | 0.0044 | 1.67E-08 | 0.001 | 0.0047 | 0.845 |
| BMR | 1:177929986:G:C | C | 0.206 | 45172 | 0.031 | 0.0053 | 2.44E-09 | 0.001 | 0.0055 | 0.869 |
| BMR | 6:7727038:G:A | A | 0.467 | 45172 | 0.024 | 0.0043 | 2.34E-08 | 0.000 | 0.0045 | 0.933 |
| BMR | 20:35437976:G:A | G | 0.407 | 45172 | 0.038 | 0.0043 | 5.94E-19 | 0.000 | 0.0045 | 0.936 |
| BMR | 20:33745375:G:T | T | 0.254 | 45165 | -0.026 | 0.0049 | 5.44E-08 | 0.000 | 0.0050 | 0.947 |
| BMR | 17:30899612:T:A | A | 0.382 | 45168 | -0.027 | 0.0044 | 6.04E-10 | 0.000 | 0.0044 | 0.956 |
| BMR | 12:906303:G:C | C | 0.197 | 45172 | 0.030 | 0.0053 | 1.62E-08 | 0.000 | 0.0057 | 0.977 |
| BMR | 8:134600502:A:G | G | 0.407 | 45172 | -0.023 | 0.0043 | 9.51E-08 | 0.000 | 0.0046 | 0.988 |
| BMR | 16:30010081:C:T | T | 0.394 | 45172 | 0.026 | 0.0043 | 1.01E-09 | 0.000 | 0.0044 | 0.993 |
| DBP | 6:28153120:G:A | A | 0.243 | 45660 | -0.043 | 0.0074 | 5.13E-09 | -0.020 | 0.0075 | 0.008 |
| DBP | 12:111446804:T:C | T | 0.482 | 45660 | 0.047 | 0.0064 | 1.82E-13 | 0.002 | 0.0066 | 0.704 |
| FEV | 4:145159425:D:1 | A | 0.359 | 41429 | 0.026 | 0.0050 | 3.43E-07 | 0.013 | 0.0051 | 0.009 |
| FEV | 6:35424010:C:T | C | 0.228 | 41430 | -0.031 | 0.0058 | 1.12E-07 | -0.003 | 0.0060 | 0.632 |
| FEV | 19:8605262:C:T | T | 0.037 | 41430 | -0.073 | 0.0130 | 2.07E-08 | -0.006 | 0.0144 | 0.677 |
| FEV | 15:83899752:A:G | A | 0.475 | 41430 | -0.026 | 0.0048 | 7.28E-08 | -0.001 | 0.0049 | 0.906 |
| FEV | 4:105897896:G:A | A | 0.257 | 41430 | -0.035 | 0.0055 | 2.28E-10 | 0.000 | 0.0056 | 0.949 |
| FIS | 1:43569801:G:A | A | 0.376 | 37351 | 0.041 | 0.0072 | 1.07E-08 | 0.036 | 0.0072 | 0.000 |
| FIS | 10:102231624:G:A | G | 0.377 | 37351 | 0.037 | 0.0072 | 3.29E-07 | 0.025 | 0.0072 | 0.000 |
| FIS | 14:32823916:A:G | A | 0.463 | 37351 | -0.037 | 0.0070 | 1.70E-07 | -0.009 | 0.0070 | 0.209 |

| | | | | | | | | | | |
|---|---|---|---|---|---|---|---|---|---|---|
| FIS | 6:28301047:A:G | G | 0.117 | 37351 | 0.061 | 0.0108 | 1.22E-08 | 0.013 | 0.0108 | 0.223 |
| FIS | 3:49805448:I:1 | TG | 0.499 | 37351 | -0.040 | 0.0070 | 6.64E-09 | -0.001 | 0.0070 | 0.894 |
| FIS | 11:64242407:G:A | NA | 0.083 | 37351 | -0.065 | 0.0126 | 2.57E-07 | NA | NA | NA |
| FVC | 2:55922340:C:G | G | 0.192 | 41430 | -0.033 | 0.0059 | 2.59E-08 | -0.010 | 0.0061 | 0.093 |
| FVC | 15:100152748:G:A | A | 0.110 | 41430 | -0.039 | 0.0074 | 1.50E-07 | -0.001 | 0.0075 | 0.910 |
| FVC | 19:8605262:C:T | T | 0.037 | 41430 | -0.081 | 0.0125 | 7.10E-11 | -0.001 | 0.0136 | 0.967 |
| hBMD | 7:121329915:I:2 | GCT | 0.261 | 7066 | 0.163 | 0.0183 | 4.37E-19 | 0.075 | 0.0194 | 0.000 |
| HC | 16:284580:G:C | C | 0.284 | 46133 | -0.038 | 0.0073 | 2.51E-07 | -0.018 | 0.0075 | 0.018 |
| HC | 15:67824962:T:A | A | 0.225 | 46133 | -0.045 | 0.0079 | 8.28E-09 | 0.002 | 0.0079 | 0.844 |
| HC | 12:882140:G:A | A | 0.201 | 46126 | 0.042 | 0.0081 | 2.53E-07 | 0.001 | 0.0083 | 0.868 |
| HC | 16:31110472:G:A | A | 0.359 | 46133 | -0.035 | 0.0068 | 3.08E-07 | -0.001 | 0.0070 | 0.893 |
| HC | 20:35437976:G:A | G | 0.407 | 46133 | 0.038 | 0.0067 | 1.09E-08 | -0.001 | 0.0067 | 0.937 |
| HGSR | 17:63842363:A:G | G | 0.331 | 45949 | -0.026 | 0.0048 | 4.81E-08 | -0.018 | 0.0048 | 0.0002 |
| HT | 22:45371575:A:G | G | 0.468 | 46116 | 0.023 | 0.0045 | 2.92E-07 | 0.018 | 0.0049 | 0.0002 |
| HT | 2:219309183:C:T | T | 0.115 | 46082 | -0.036 | 0.0070 | 2.01E-07 | -0.029 | 0.0078 | 0.0003 |
| HT | 8:134637605:G:A | A | 0.259 | 46116 | -0.029 | 0.0052 | 2.64E-08 | -0.021 | 0.0058 | 0.0003 |
| HT | 5:32711527:C:A | A | 0.195 | 46116 | -0.034 | 0.0057 | 1.57E-09 | -0.021 | 0.0060 | 0.0005 |
| HT | 2:23958289:G:T | T | 0.188 | 46032 | -0.033 | 0.0058 | 1.14E-08 | -0.021 | 0.0063 | 0.0007 |
| HT | 5:132336076:T:C | T | 0.299 | 46112 | -0.026 | 0.0048 | 7.51E-08 | -0.015 | 0.0051 | 0.0037 |
| HT | 10:103076290:T:C | C | 0.389 | 46116 | 0.024 | 0.0046 | 2.71E-07 | 0.013 | 0.0050 | 0.0068 |
| HT | 18:23135692:G:A | A | 0.495 | 46115 | 0.031 | 0.0045 | 9.28E-12 | 0.012 | 0.0046 | 0.0091 |
| HT | 11:65965994:G:A | A | 0.061 | 46116 | -0.069 | 0.0095 | 4.00E-13 | -0.026 | 0.0103 | 0.011 |
| HT | 5:108820558:T:A | A | 0.228 | 46116 | 0.029 | 0.0054 | 6.06E-08 | 0.014 | 0.0054 | 0.011 |
| HT | 1:16986959:G:A | A | 0.234 | 46116 | -0.035 | 0.0053 | 5.34E-11 | -0.014 | 0.0063 | 0.026 |
| HT | 1:47333967:G:C | G | 0.499 | 46116 | -0.023 | 0.0045 | 2.21E-07 | -0.011 | 0.0049 | 0.029 |
| HT | 2:232210371:C:T | T | 0.028 | 46116 | -0.079 | 0.0137 | 6.67E-09 | -0.029 | 0.0157 | 0.062 |
| HT | 17:7455201:C:T | T | 0.208 | 46100 | 0.031 | 0.0055 | 1.34E-08 | 0.010 | 0.0058 | 0.098 |
| HT | 15:61910283:C:T | C | 0.457 | 46116 | 0.024 | 0.0045 | 1.18E-07 | 0.008 | 0.0048 | 0.114 |
| HT | 9:95447312:G:A | A | 0.338 | 46116 | -0.030 | 0.0048 | 2.23E-10 | -0.008 | 0.0057 | 0.143 |
| HT | 6:151807942:T:C | C | 0.477 | 46116 | 0.027 | 0.0045 | 3.70E-09 | 0.007 | 0.0048 | 0.168 |
| HT | 3:53103295:C:G | C | 0.402 | 46116 | 0.028 | 0.0046 | 9.44E-10 | 0.006 | 0.0051 | 0.269 |
| HT | 6:34871867:G:A | A | 0.016 | 46116 | -0.107 | 0.0182 | 3.79E-09 | -0.018 | 0.0204 | 0.383 |
| HT | 6:19838216:C:A | A | 0.062 | 46116 | 0.076 | 0.0094 | 5.89E-16 | -0.008 | 0.0100 | 0.396 |
| HT | 3:141608632:G:A | A | 0.395 | 46116 | 0.033 | 0.0046 | 6.93E-13 | 0.004 | 0.0048 | 0.430 |
| HT | 9:96388497:A:G | G | 0.163 | 46116 | 0.036 | 0.0061 | 4.99E-09 | 0.005 | 0.0071 | 0.476 |
| HT | 12:102195300:D:2 | T | 0.011 | 46101 | -0.111 | 0.0212 | 1.73E-07 | -0.015 | 0.0229 | 0.513 |
| HT | 17:30784350:A:G | G | 0.177 | 46116 | -0.040 | 0.0059 | 1.30E-11 | -0.004 | 0.0062 | 0.518 |
| HT | 7:66286473:T:A | A | 0.191 | 46106 | 0.030 | 0.0058 | 1.63E-07 | 0.004 | 0.0062 | 0.526 |
| HT | 4:17883363:T:C | C | 0.136 | 46116 | -0.053 | 0.0066 | 5.75E-16 | -0.004 | 0.0069 | 0.530 |
| HT | 4:144658755:G:A | A | 0.484 | 46116 | 0.028 | 0.0045 | 2.50E-10 | -0.003 | 0.0046 | 0.560 |
| HT | 20:49158557:C:T | T | 0.238 | 46116 | 0.029 | 0.0053 | 6.82E-08 | 0.003 | 0.0057 | 0.631 |
| HT | 1:88770346:T:A | T | 0.457 | 46116 | -0.026 | 0.0045 | 1.44E-08 | -0.002 | 0.0046 | 0.656 |
| HT | 2:55922340:C:G | G | 0.192 | 46116 | -0.039 | 0.0057 | 1.12E-11 | 0.003 | 0.0061 | 0.662 |
| HT | 13:49663119:G:A | A | 0.022 | 46116 | 0.082 | 0.0153 | 7.45E-08 | -0.005 | 0.0158 | 0.732 |
| HT | 8:129748784:G:C | C | 0.464 | 46116 | -0.023 | 0.0045 | 1.88E-07 | -0.002 | 0.0049 | 0.747 |
| HT | 6:130060101:T:C | T | 0.311 | 46116 | 0.034 | 0.0049 | 1.95E-12 | 0.002 | 0.0053 | 0.770 |
| HT | 7:92618019:A:G | G | 0.250 | 46116 | 0.047 | 0.0052 | 2.71E-19 | 0.001 | 0.0054 | 0.820 |
| HT | 10:68199640:G:A | A | 0.486 | 46103 | 0.025 | 0.0044 | 3.05E-08 | 0.001 | 0.0047 | 0.834 |
| HT | 2:25240614:G:A | A | 0.411 | 46116 | 0.031 | 0.0046 | 2.20E-11 | 0.001 | 0.0050 | 0.846 |
| HT | 14:65075889:C:T | T | 0.421 | 46115 | -0.025 | 0.0046 | 4.42E-08 | -0.001 | 0.0050 | 0.849 |
| HT | 6:34246545:C:G | C | 0.090 | 46116 | 0.063 | 0.0078 | 7.82E-16 | 0.002 | 0.0089 | 0.853 |
| HT | 1:41075189:T:C | T | 0.219 | 46115 | 0.035 | 0.0055 | 1.83E-10 | 0.001 | 0.0058 | 0.869 |
| HT | 6:34857885:T:C | C | 0.138 | 46116 | 0.041 | 0.0065 | 2.88E-10 | -0.001 | 0.0074 | 0.887 |
| HT | 17:63830956:G:A | A | 0.284 | 46116 | 0.036 | 0.0050 | 4.18E-13 | 0.001 | 0.0061 | 0.891 |
| HT | 2:88575373:C:A | C | 0.281 | 46116 | 0.030 | 0.0050 | 2.67E-09 | -0.001 | 0.0051 | 0.907 |
| HT | 1:149934520:T:C | C | 0.407 | 46116 | 0.040 | 0.0046 | 2.05E-18 | 0.001 | 0.0047 | 0.912 |
| HT | 12:123341012:C:T | T | 0.203 | 46116 | 0.032 | 0.0056 | 1.07E-08 | -0.001 | 0.0058 | 0.913 |
| HT | 15:100152748:G:A | A | 0.110 | 46116 | -0.058 | 0.0072 | 6.73E-16 | -0.001 | 0.0084 | 0.919 |
| HT | 19:4954443:G:A | A | 0.198 | 46116 | -0.030 | 0.0056 | 1.01E-07 | 0.001 | 0.0064 | 0.921 |
| HT | 15:67165360:A:G | G | 0.056 | 46116 | 0.069 | 0.0097 | 2.00E-12 | -0.001 | 0.0106 | 0.929 |
| HT | 6:7727038:G:A | A | 0.466 | 46116 | 0.033 | 0.0045 | 3.82E-13 | 0.000 | 0.0048 | 0.945 |
| HT | 20:33745375:G:T | T | 0.254 | 46109 | -0.036 | 0.0052 | 5.97E-12 | 0.000 | 0.0056 | 0.946 |
| HT | 15:83913152:T:C | T | 0.476 | 46116 | -0.036 | 0.0045 | 1.22E-15 | 0.000 | 0.0051 | 0.951 |
| HT | 17:29562968:T:C | T | 0.343 | 46116 | -0.029 | 0.0047 | 9.39E-10 | 0.000 | 0.0050 | 0.951 |
| HT | 1:184051811:G:A | A | 0.348 | 46116 | 0.032 | 0.0048 | 2.89E-11 | 0.000 | 0.0049 | 0.951 |
| HT | 7:2762888:T:C | C | 0.301 | 46116 | -0.039 | 0.0049 | 4.65E-15 | 0.000 | 0.0051 | 0.952 |
| HT | 6:142403781:C:T | T | 0.279 | 46115 | -0.045 | 0.0050 | 4.44E-19 | 0.000 | 0.0054 | 0.954 |
| HT | 3:172447937:C:T | T | 0.313 | 46116 | 0.025 | 0.0049 | 2.58E-07 | 0.000 | 0.0050 | 0.957 |
| HT | 3:129301935:A:G | A | 0.219 | 46115 | -0.030 | 0.0055 | 5.06E-08 | 0.000 | 0.0058 | 0.968 |
| HT | 19:55482069:G:T | T | 0.026 | 46116 | -0.073 | 0.0143 | 3.53E-07 | -0.001 | 0.0154 | 0.972 |

| | | | | | | | | | | |
|---|---|---|---|---|---|---|---|---|---|---|
| HT | 6:26183874:G:A | A | 0.258 | 46116 | -0.044 | 0.0052 | 3.47E-17 | 0.000 | 0.0058 | 0.980 |
| HT | 5:177089630:G:A | A | 0.242 | 46116 | 0.042 | 0.0053 | 2.99E-15 | 0.000 | 0.0056 | 0.983 |
| HT | 12:28484182:G:A | A | 0.316 | 46115 | -0.035 | 0.0048 | 3.53E-13 | 0.000 | 0.0052 | 0.983 |
| HT | 20:35437976:G:A | G | 0.407 | 46116 | 0.059 | 0.0046 | 1.94E-37 | 0.000 | 0.0050 | 0.987 |
| HT | 19:8605262:C:T | T | 0.037 | 46116 | -0.100 | 0.0120 | 1.01E-16 | 0.000 | 0.0149 | 0.988 |
| HT | 11:75566583:A:G | A | 0.115 | 46116 | 0.040 | 0.0071 | 1.53E-08 | 0.000 | 0.0078 | 0.994 |
| HT | 15:88857449:A:G | G | 0.029 | 46116 | -0.120 | 0.0135 | 8.12E-19 | 0.000 | 0.0160 | 0.998 |
| MTCIM | 5:175510190:C:T | T | 0.133 | 46025 | -0.048 | 0.0091 | 1.10E-07 | NA | NA | NA |
| PEF | 9:133370796:C:A | A | 0.137 | 41339 | -0.044 | 0.0078 | 2.18E-08 | -0.027 | 0.0078 | 0.0006 |
| PR | 20:44311053:C:T | T | 0.015 | 37166 | 0.163 | 0.0297 | 3.86E-08 | 0.163 | 0.0298 | 4.68E-08 |
| PR | 14:23396676:G:A | A | 0.357 | 37166 | 0.070 | 0.0076 | 5.34E-20 | 0.003 | 0.0076 | 0.711 |
| PR | 20:38213354:T:C | C | 0.474 | 37166 | -0.074 | 0.0073 | 1.86E-24 | 0.000 | 0.0073 | 0.948 |
| PR | 7:100886760:T:G | G | 0.183 | 37166 | 0.052 | 0.0094 | 3.25E-08 | -0.001 | 0.0094 | 0.957 |
| PR | 2:178856319:G:A | A | 0.086 | 37166 | 0.094 | 0.0130 | 4.23E-13 | -0.001 | 0.0130 | 0.958 |
| SBP | 5:32713221:T:C | C | 0.393 | 45659 | -0.035 | 0.0061 | 1.20E-08 | -0.013 | 0.0063 | 0.044 |
| SBP | 11:47792728:A:G | G | 0.454 | 45659 | 0.031 | 0.0060 | 2.62E-07 | 0.007 | 0.0061 | 0.267 |
| SBP | 16:24823847:G:A | A | 0.194 | 45659 | -0.044 | 0.0076 | 6.16E-09 | 0.001 | 0.0076 | 0.934 |
| SBP | 1:11823674:C:T | T | 0.162 | 45659 | -0.049 | 0.0081 | 2.35E-09 | 0.000 | 0.0085 | 0.969 |
| WT | 9:108990879:G:T | T | 0.075 | 46067 | 0.061 | 0.0109 | 2.05E-08 | 0.046 | 0.0111 | 0.00003 |
| WT | 16:284580:G:C | C | 0.284 | 46067 | -0.033 | 0.0064 | 1.91E-07 | -0.015 | 0.0065 | 0.020 |
| WT | 16:30011243:A:G | G | 0.457 | 46063 | 0.031 | 0.0057 | 6.35E-08 | 0.011 | 0.0058 | 0.048 |
| WT | 15:67824962:T:A | A | 0.225 | 46067 | -0.035 | 0.0068 | 2.81E-07 | -0.012 | 0.0070 | 0.093 |
| WT | 20:35434589:C:A | C | 0.346 | 46067 | 0.041 | 0.0060 | 1.63E-11 | 0.006 | 0.0061 | 0.308 |
| WT | 17:30899612:T:A | A | 0.382 | 46063 | -0.032 | 0.0059 | 4.85E-08 | -0.003 | 0.0059 | 0.589 |
| WT | 4:145159425:D:1 | A | 0.359 | 46066 | 0.031 | 0.0060 | 2.24E-07 | 0.003 | 0.0061 | 0.630 |
| WT | 12:882140:G:A | A | 0.201 | 46060 | 0.038 | 0.0071 | 1.08E-07 | 0.001 | 0.0072 | 0.877 |
| WT | 6:130060101:T:C | T | 0.311 | 46067 | 0.032 | 0.0062 | 3.16E-07 | 0.001 | 0.0064 | 0.922 |
| WT | 1:177929986:G:C | C | 0.206 | 46067 | 0.042 | 0.0071 | 2.29E-09 | 0.000 | 0.0072 | 0.947 |
| WT | 11:27700751:G:T | T | 0.319 | 46067 | 0.033 | 0.0061 | 7.79E-08 | 0.000 | 0.0063 | 0.955 |
| WT | 18:60372043:C:T | T | 0.021 | 46067 | -0.104 | 0.0202 | 2.54E-07 | 0.000 | 0.0204 | 0.999 |
| AMeno | 1:38895943:G:A | A | 0.457 | 14083 | -0.064 | 0.0113 | 1.44E-08 | -0.028 | 0.0114 | 0.015 |
| AMeno | 19:55319820:A:G | G | 0.391 | 14088 | -0.084 | 0.0118 | 8.53E-13 | -0.009 | 0.0119 | 0.441 |
| AMeno | 4:83446133:G:A | A | 0.476 | 14075 | 0.061 | 0.0111 | 4.61E-08 | 0.004 | 0.0111 | 0.710 |
| AMeno | 6:10887043:C:G | C | 0.174 | 14088 | 0.078 | 0.0150 | 1.80E-07 | 0.002 | 0.0156 | 0.911 |
| AMeno | 12:66310445:A:G | G | 0.031 | 14088 | 0.214 | 0.0331 | 8.82E-11 | 0.002 | 0.0333 | 0.960 |
| AMeno | 8:38030499:C:G | C | 0.225 | 14087 | 0.084 | 0.0136 | 6.66E-10 | 0.000 | 0.0136 | 0.974 |
| AMeno | 20:5967581:G:A | A | 0.061 | 14088 | 0.232 | 0.0238 | 1.35E-22 | 0.000 | 0.0238 | 1.000 |
| BFP | 20:63738996:T:C | T | 0.340 | 45131 | -0.027 | 0.0051 | 9.07E-08 | -0.003 | 0.0051 | 0.574 |

**Supplementary Figures**

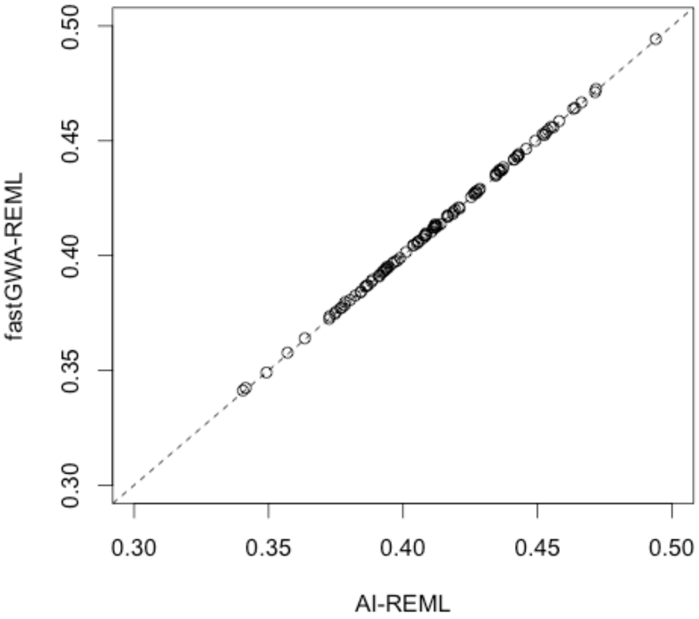


**Supplementary Figure 1.** Comparison between fastGWA-REML and AI-REML. The phenotypes were simulated based on real genotypes of 100,000 individuals from the UKB with $V_g = 0.4$ (see **Supplementary Note 5** for details of the simulation method and data). Plotted are the $\hat{\sigma}_g^2$ estimated by fastGWA-REML against those estimated by the AI-REML in GCTA. Each dot represents one simulation replicate (100 simulations in total). The correlation of $\hat{\sigma}_g^2$ between the two methods is > 0.9999.

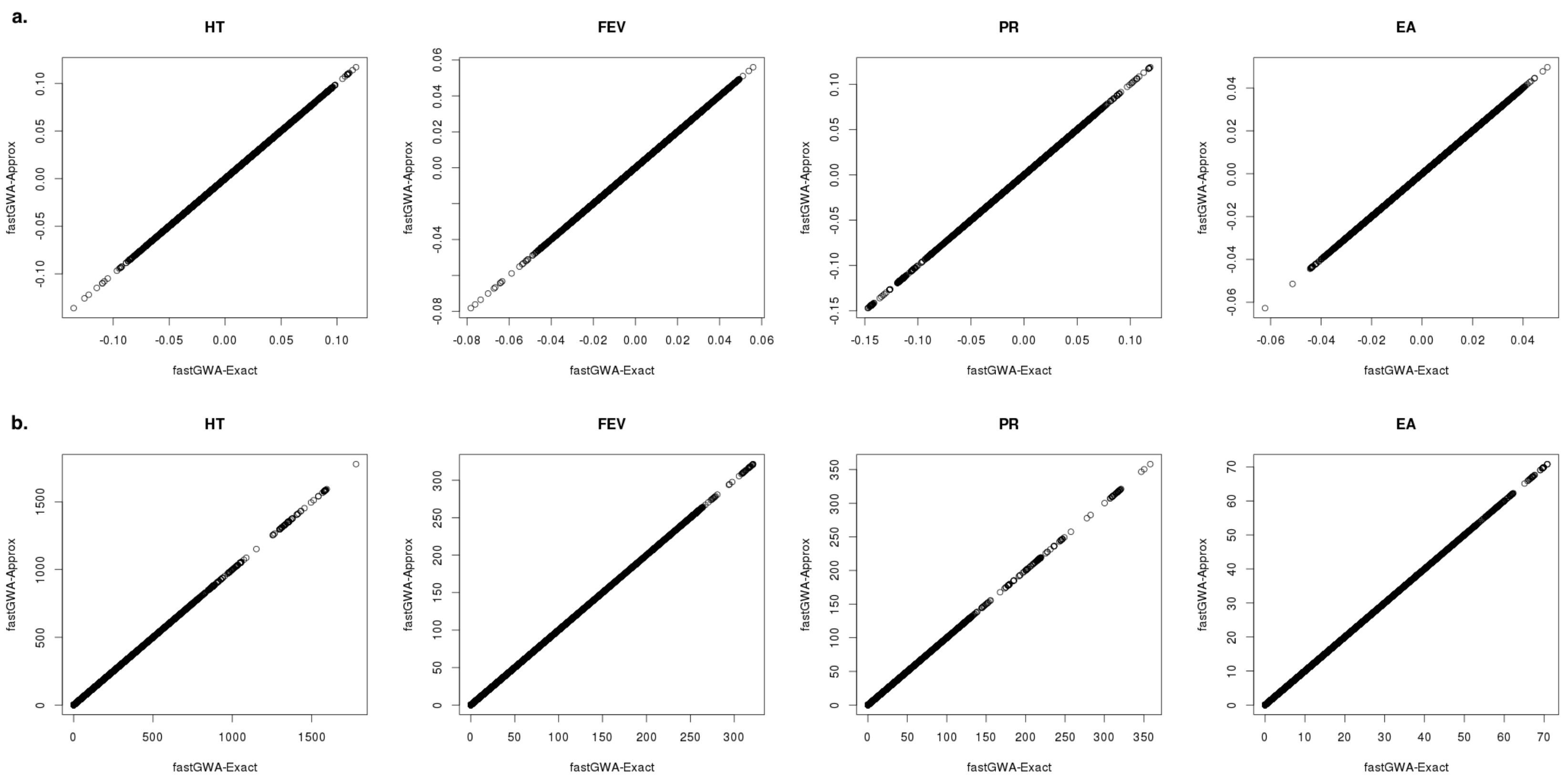


**Supplementary Figure 2.** Comparison between the approximate and exact fastGWA tests. We selected four quantitative traits from the UKB for comparison, including height (HT, $n_{\mathrm{HT}}$ = 455,332), forced expiratory volume in 1-second (FEV, $n_{\mathrm{FEV}}$ = 415,931), pulse rate (PR, $n_{\mathrm{PR}}$ = 149,082), and education attainment (EA, $n_{\mathrm{EA}}$ = 304,998) (see **Supplementary Table 1** for more information about the traits). Plotted are the estimated variant effects (in Panel a.) or $\chi^2$-statistics (in Panel b.) of 8,531,416 variants computed by the exact fastGWA method (fastGWA-Exact) against those by the fastGWA test using the GRAMMAR-GAMMA approximation (see **Supplementary Note 2** for details). The correlation of the estimated variant effect or $\chi^2$-statistic between the two methods is > 0.9999 for all the four traits.

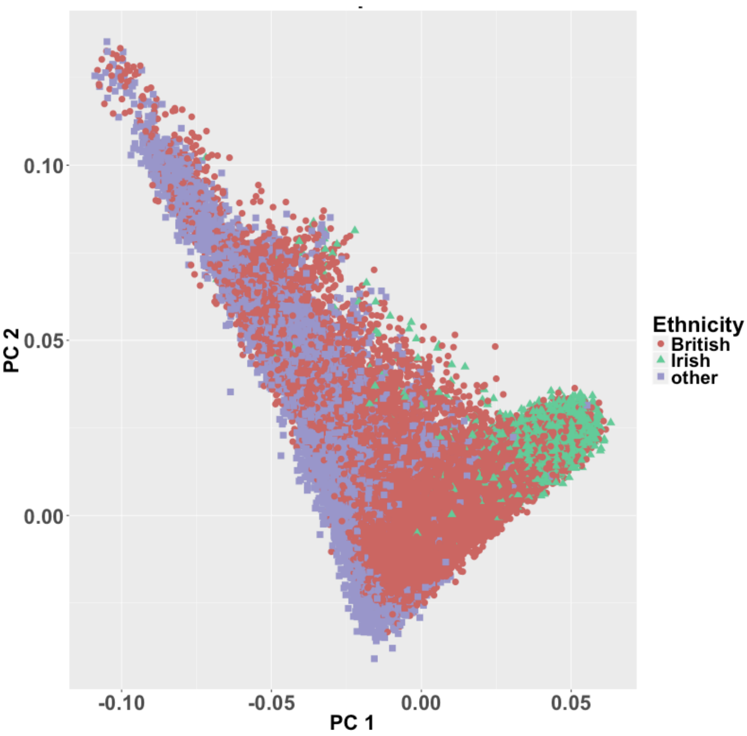


**Supplementary Figure 3.** The first and second principal components (PC1 and PC2) of all the UKB participants of European ancestry compared to their self-reported ethnicity. The red dots represent the ones self-reported as "British", the green dots represent those self-reported as "Irish", and the purple dots represent those self-reported as "other-white background".

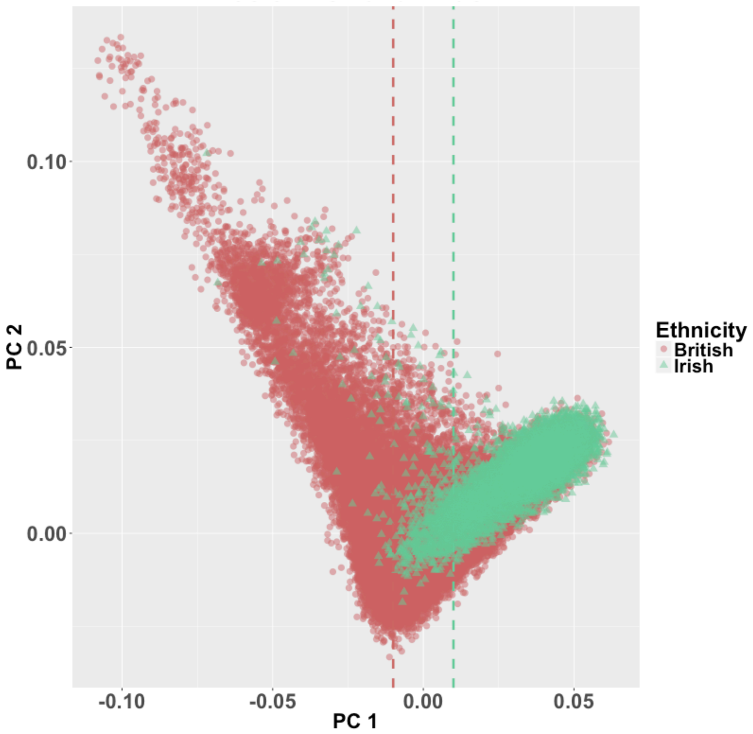


**Supplementary Figure 4.** The first and second principal components (PC1 and PC2) plotted against self-reported ethnicity among individuals of Irish and British ancestry from the UKB. In the simulation, we randomly selected 9,000 "Irish" individuals from the green dots on the right-hand side of the green vertical line (PC1 ≥ 0.01), and 9,000 "British" individuals from the red dots on the left-hand side of the red vertical line (PC1 ≤ -0.01) (see **Supplementary Note 4** for details of the simulation).

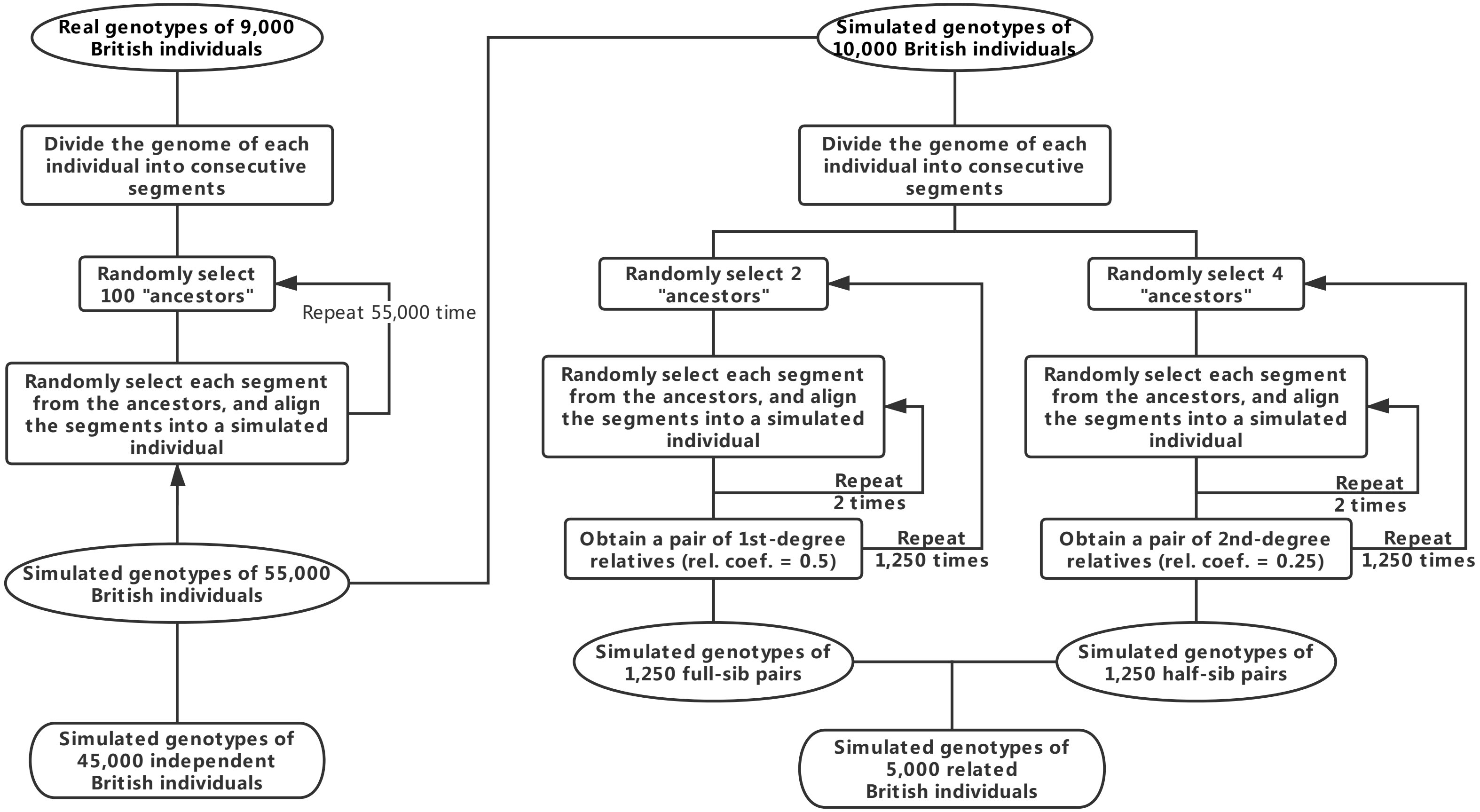


**Supplementary Figure 5.** Schematic diagram of simulating a GWAS data set with relatedness and population stratification from existing GWAS data. "rel. coef.": relatedness coefficient.

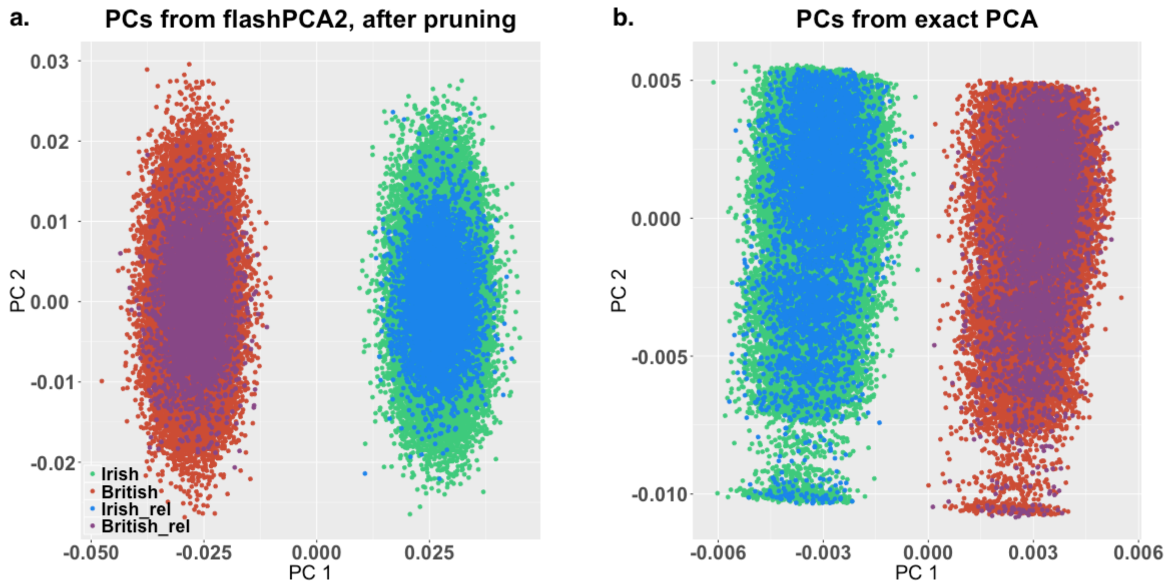


**Supplementary Figure 6**. Principal component analysis of all the 100,000 simulated individuals. The left panel (a) shows the first two PCs (PC1 and PC2) computed from a set of pruned variants (window size = 1 Mb, step size =50 and LD $r^2$ threshold = 0.05) by flashPCA2, and the right panel (b) shows the first two PCs from exact PCA without LD pruning implemented in GCTA. The red dots represent the simulated British individuals, and the green dots represent the simulated Irish individuals. In panels a) and b), the related individuals (relatedness coefficients ≥ 0.05) were labelled with a slightly darker colour in each group (“Irish_rel” and “British_rel”).

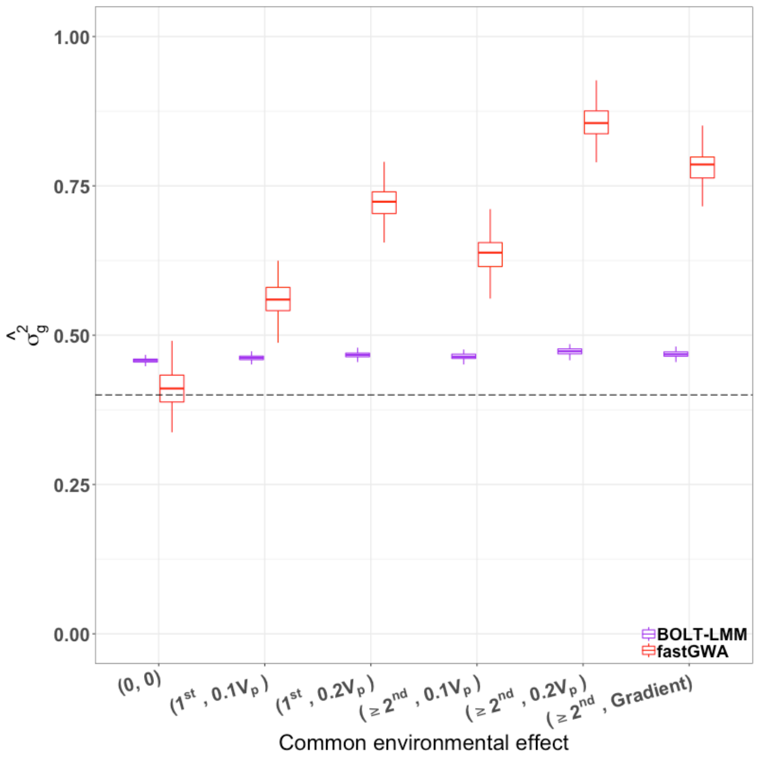


**Supplementary Figure 7.** Comparison of $\hat{\sigma}_g^2$ estimated by fastGWA-REML to that by BOLT-REML (used in BOLT-LMM) at different degrees of relatedness in simulations. The x-axis represents different degrees of relatedness, where (0, 0) represents no common environmental effect; ($1^{st}$, $0.1V_p$) or ($1^{st}$, $0.2V_p$) represents that common environmental effects explained 10% or 20% of the phenotypic variance ($V_p$) among $1^{st}$ degree relatives; (≥$2^{st}$, $0.1V_p$) or (≥$2^{st}$, $0.2V_p$) represents that common environmental effects explained 10% or 20% of $V_p$ among all pairs of the $1^{st}$ and $2^{nd}$ degree relatives; (≥$2^{st}$, Gradient) represents that common environmental effects explained 20% of $V_p$ among the $1^{st}$ degree relatives and 10% of $V_p$ among the $2^{nd}$ degree relatives. The y-axis represents the value of $\hat{\sigma}_g^2$. The black dashed line represents the true simulation parameter ($h^2$ = 0.4).

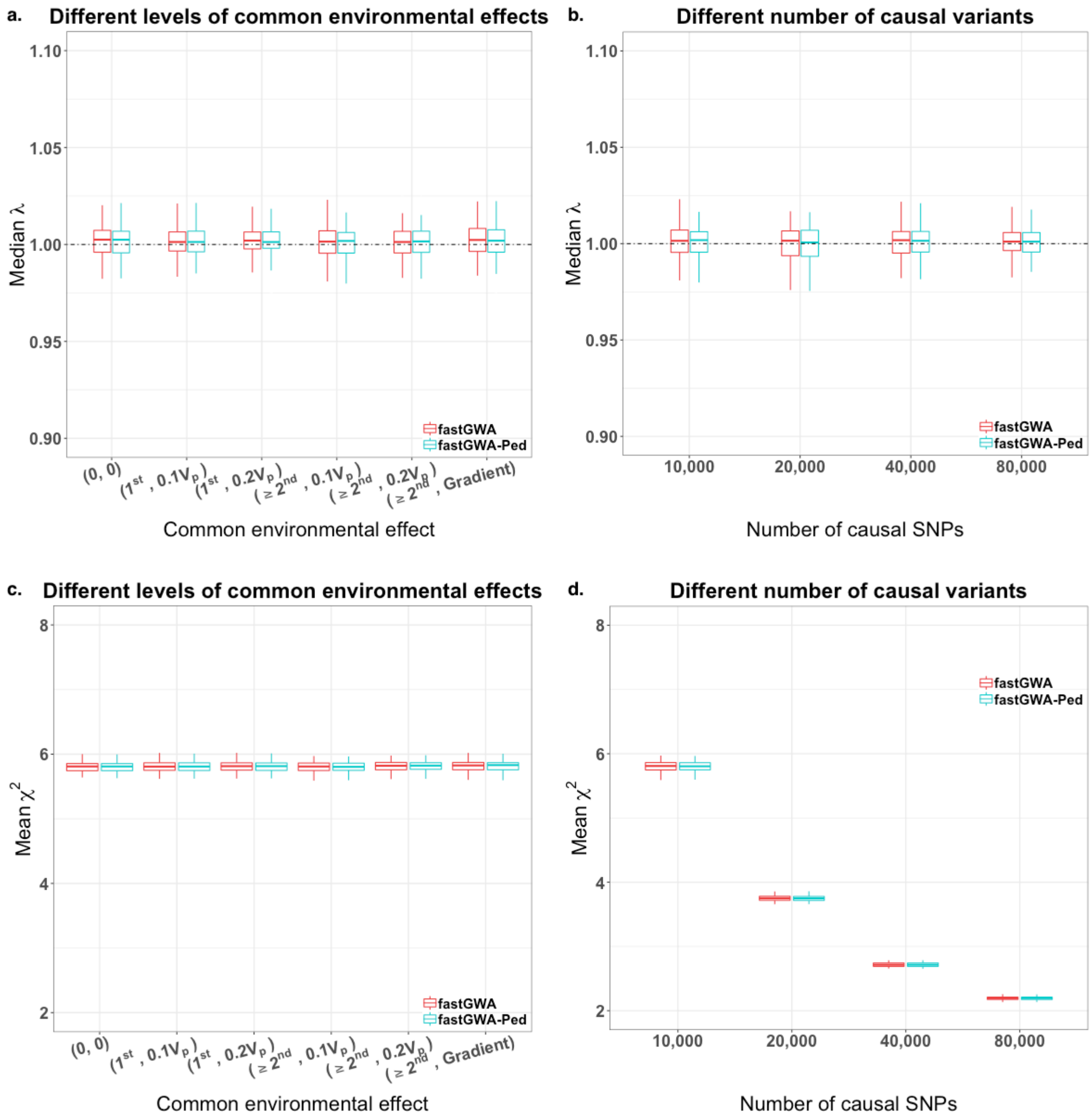


**Supplementary Figure 8.** Comparison between fastGWA and fastGWA-Ped. Panels a) and b) show the median λ at the null variants for fastGWA and fastGWA-Ped, respectively. Panel a) shows the median λ with different levels of common environmental effects, and panel b) shows the median λ with different number of simulated causal variants. Panels c) and d) show the mean $\chi^2$ value of causal variants for fastGWA and fastGWA-Ped. Panel c) shows the mean $\chi^2$ value with different levels of common environmental effects, and panel d) shows the mean $\chi^2$ value with different number of simulated causal variants. In all the panels, each box plot represents the distribution of the estimates (i.e., median λ and mean $\chi^2$) across 100 simulation replicates.

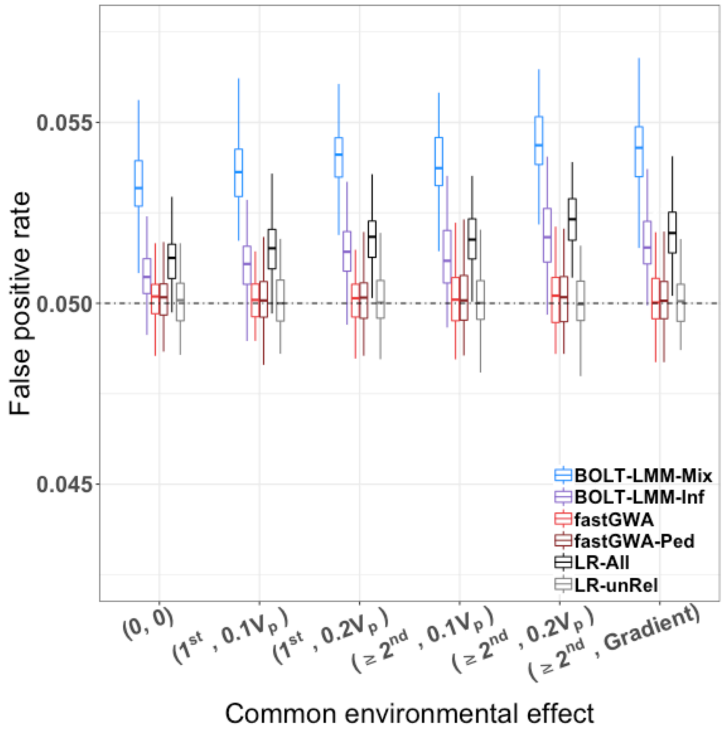


**Supplementary Figure 9.** Comparison of false positive rate (FPR) among different association methods. We used the simulated data as presented in **Figures 1** and **2** to compute the FPR of each association method across different simulation scenarios with different levels of common environmental effects. Each boxplot represents the distribution of FPR across 100 simulation replicates.

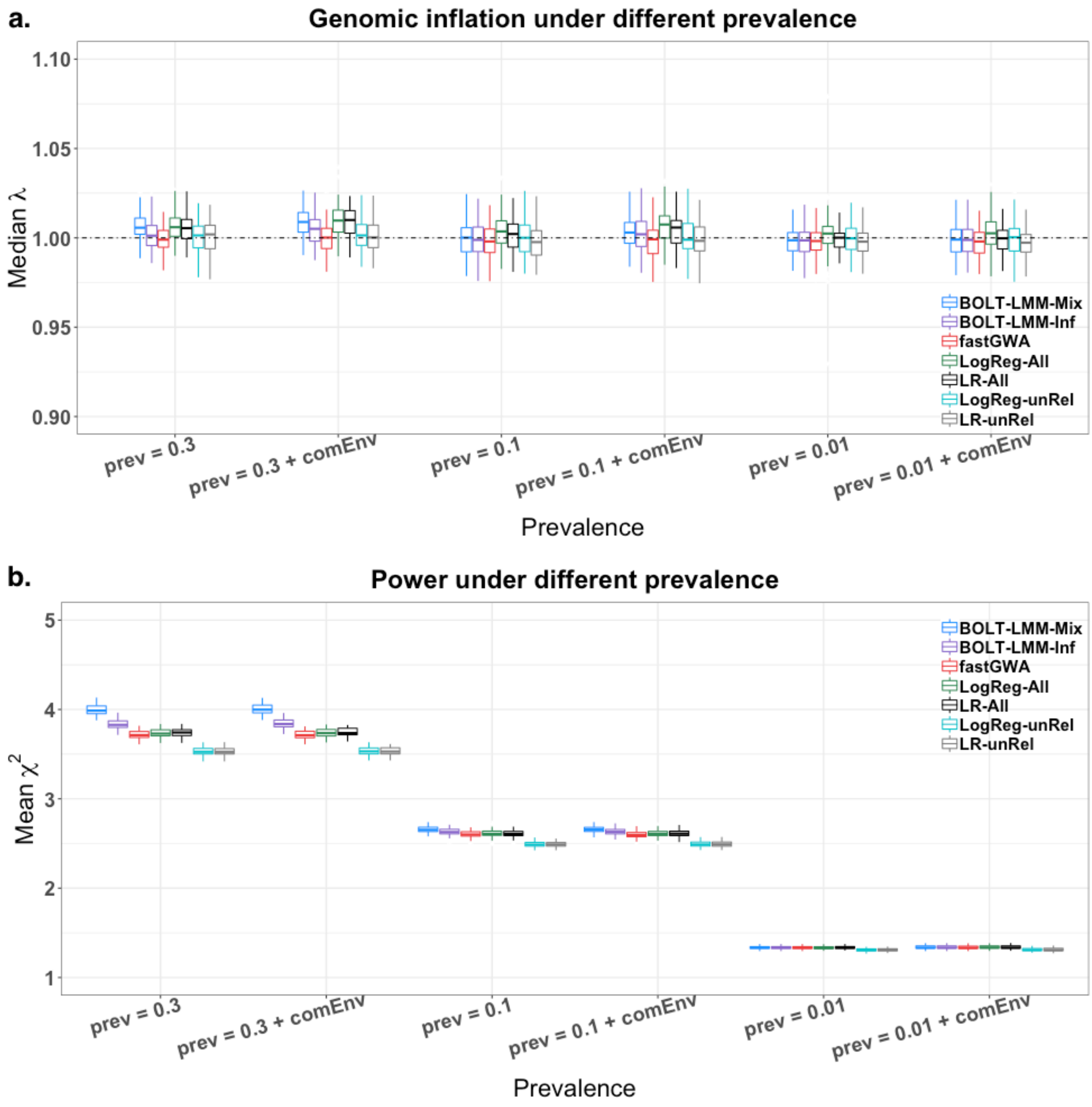


**Supplementary Figure 10.** Comparison of genomic inflation factor and power among different association methods for binary phenotypes in unbalanced case-control settings. We simulated the binary phenotypes using the method described in **Supplementary Note 5** with sample prevalence rates of 0.3, 0.1, and 0.01. The common environmental effect (labelled with "+ comEnv") was generated using the same setting as in the ($\geq 2^{nd}$, $0.1V_p$) case shown in **Figure 1**. We performed logistic regression and linear regression analyses in all individuals (LogReg-All and LR-All, respectively) and unrelated individuals (LogReg-unRel and LR-unRel, respectively), using PLINK2. We also performed BOLT-LMM-Mix and BOLT-LMM-Inf in all individuals. We then compared the results from fastGWA against these methods. We computed the genomic inflation factor at the null variants and the statistical power at the causal variants.

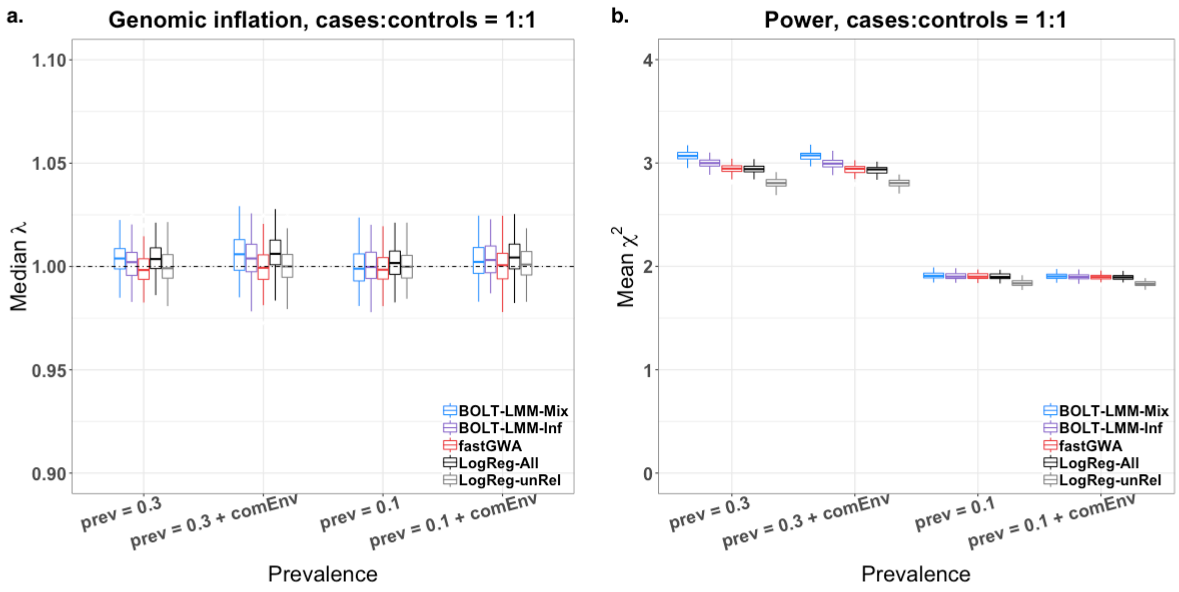


**Supplementary Figure 11.** Comparison of genomic inflation and power among different association methods for binary phenotypes in balanced case-control settings. Using the data generated by the simulations presented in **Supplementary Figure 8**, we randomly down-sampled the controls to perform a balanced case-control analysis. We ignored the case with prevalence rate of 0.01 because the number of cases is too small (only ~1000 cases). There were ~30,000 case and ~30,000 controls in the scenario with prevalence of 0.3, and ~10,000 case and ~10,000 controls in the scenario with prevalence of 0.1. We then performed the same association analyses as shown in **Supplementary Figure 8** to compare the methods.

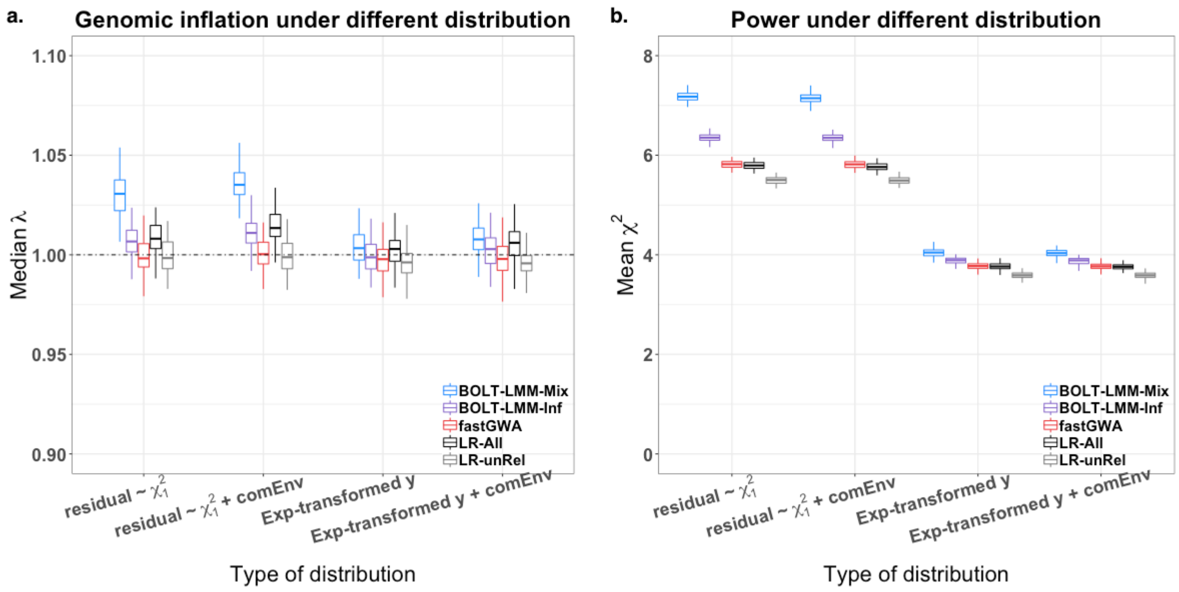


**Supplementary Figure 12.** Comparison of genomic inflation and power among different association methods for non-normally distributed phenotypes. Two classes of non-normally distributed phenotypes were investigated, including phenotypes with residuals following a $\chi_1^2$ distribution, and phenotypes being exponentially transformed (see **Supplementary Note 5** for details of the simulations). The common environmental effect (labelled with "+ comEnv") was generated using the same setting as in the ($\geq 2^{nd}$, $0.1V_p$) case shown in **Figure 1**.

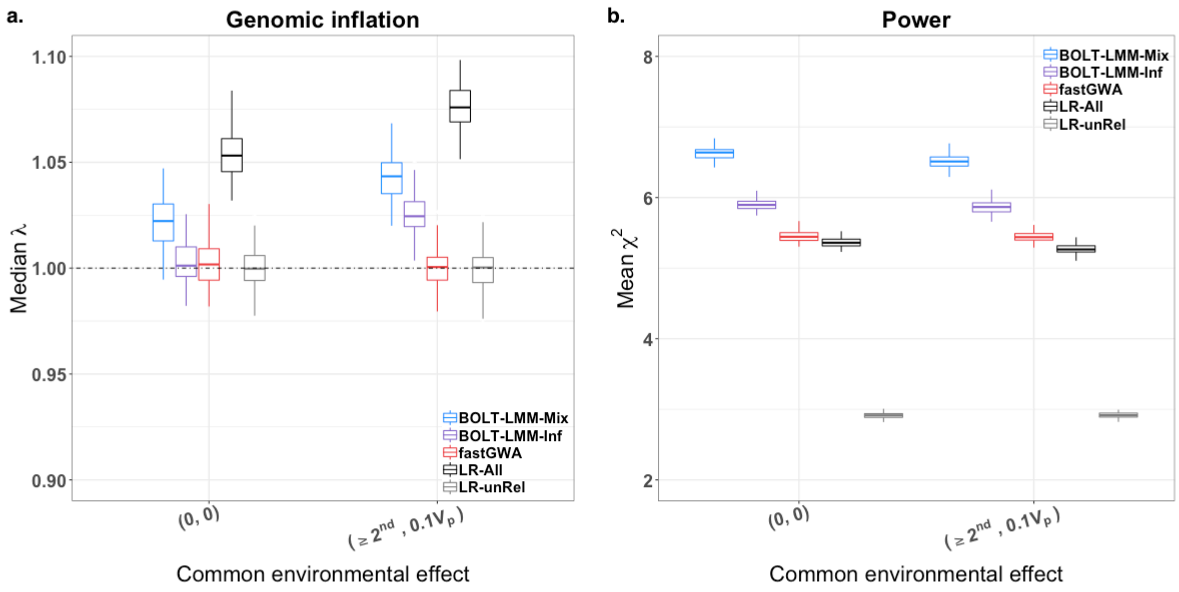


**Supplementary Figure 13.** Comparison of genomic inflation and power among different association methods for phenotypes simulated based on real genotypes from the UKB. We extracted SNP array genotyped data of 556,042 variants on 100,000 individuals of European ancestry from the UKB. The individuals were purposely sampled to achieve a higher sparse GRM density (85,727 unique pairs of related individuals; sparse GRM density = $2.7 \times 10^{-5}$ ) than that in the whole UKB sample (sparse GRM density =$3.9\times 10^{-6}$). Two simulation settings were examined, including phenotypes simulated without any common environmental effect (same as the (0, 0) case in **Figure 1**), and phenotypes simulated with common environmental effect among the first- and second-degree relatives (same as the ($\geq 2^{nd}$, $0.1V_p$) case in **Figure 1**). The difference in power between LR-unRel and the other methods was due to a large proportion of related individuals being excluded ($N_{LR\text{-}unRel}$ = 41,701).

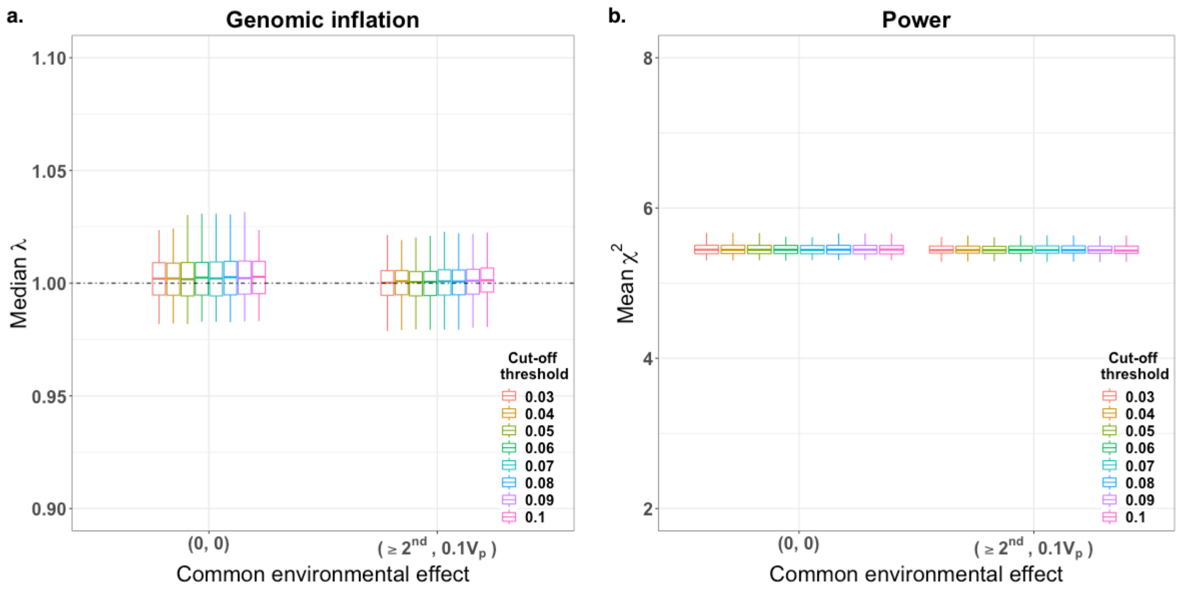


**Supplementary Figure 14.** Genomic inflation and power of fastGWA with the sparse GRM thresholded at different genetic relatedness cut-off values. This simulation was performed based on real genotypes from the UKB (see simulation settings in **Supplementary Note 5**). We constructed different sparse GRMs by setting off-diagonal elements below a certain threshold (varying from 0.03 to 0.10) to zero and performed fastGWA analyses using these sparse GRMs.

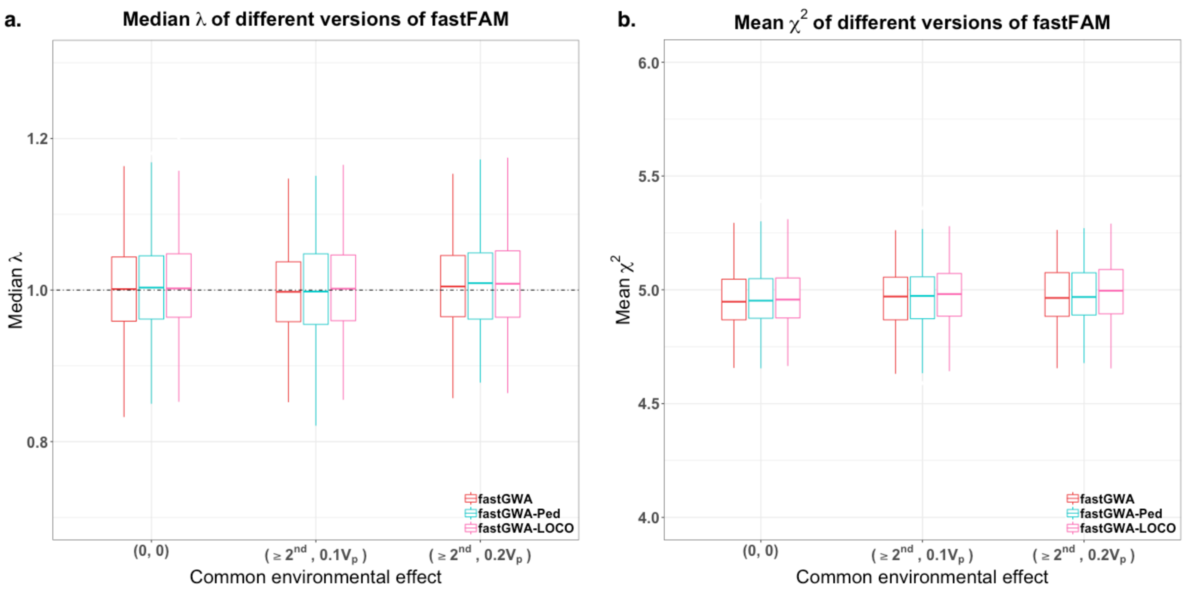


**Supplementary Figure 15.** Comparison of genomic inflation and power between fastGWA, fastGWA-LOCO, and fastGWA-Ped. Shown are the results from the analyses of a simulated data set based on the simulation strategy described in **Supplementary Note 5** (with $\sigma_g^2 = 0.4V_p$, $\sigma_c^2 = 0.1V_p, or\ 0.2V_p$ for all 1st and 2nd relatives and $\sigma_c^2 = 0$ for all unrelated individuals).

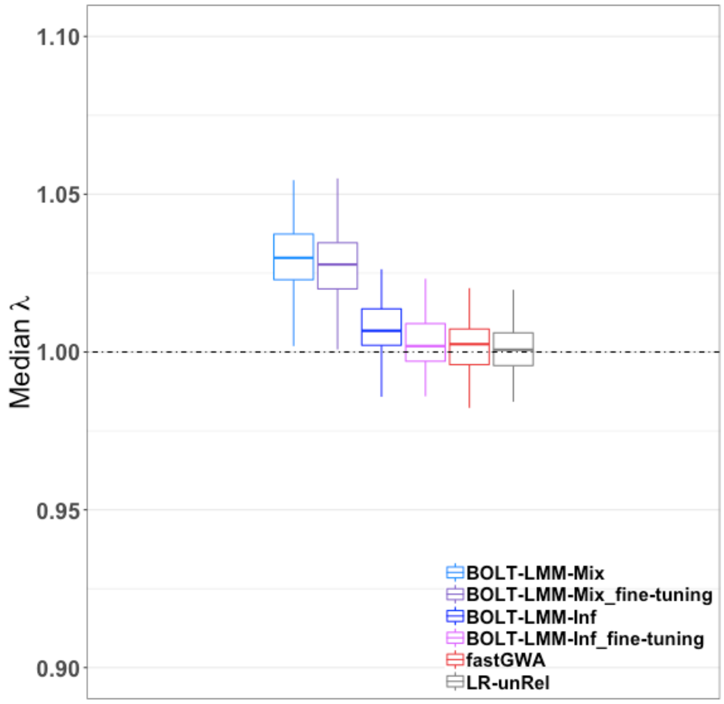


**Supplementary Figure 16.** Comparison of genomic inflation between BOLT-LMM (estimating the variance components only once using all variants) and BOLT-LMM_fine-tuning (re-estimating the variance components when a chromosome is left out). The simulation setting was the same as the (0, 0) scenario in **Figure 1**. The genomic inflation factor was computed at the null variants.

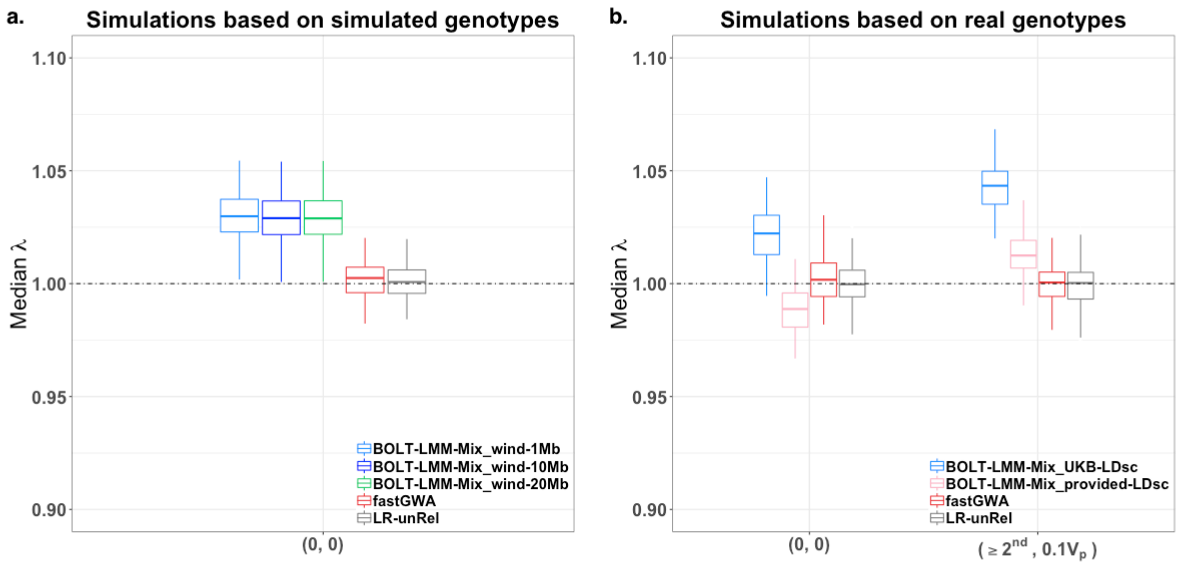


**Supplementary Figure 17.** Genomic inflation of the BOLT-LMM mixture model (BOLT-LMM-Mix) using different LD references. Shown in panel a) are the results from simulations based on the simulated genotype data (**Supplementary Note 5**) using the same setting as in the (0,0) case in **Figure 1**. The LD scores were computed from the sample using three window sizes, i.e., 1 Mb (BOLT-LMM-Mix_wind-1Mb), 10 Mb (BOLT-LMM-Mix_wind-10Mb), and 20 Mb (BOLT-LMM-Mix_wind-20Mb). Shown in panel b) are the results from simulations based on real genotypes (**Supplementary Note 5**) using the same settings as in the (0,0) and ($\geq 2^{nd}$, $0.1V_p$) cases in **Figure 1**. Two sets of LD score were tested: LD scores computed from the sample using a window size of # (BOLT-LMM-Mix_UKB-LDsc) and LD scores obtained from the BOLT-LMM website (BOLT-LMM-Mix_provided-LDsc).

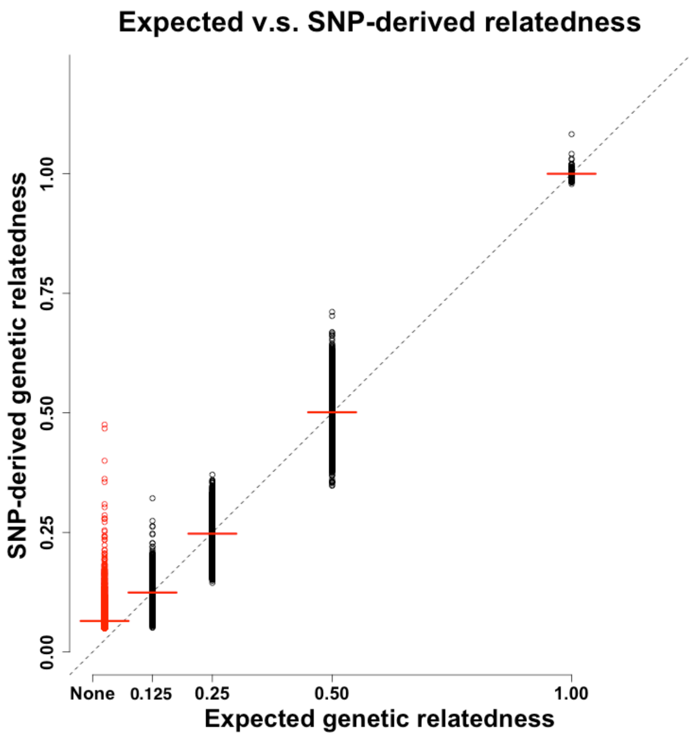


**Supplementary Figure 18.** Comparison between the reported genetic relatedness and the SNP-derived genetic relatedness of the UKB participants. The y-axis represents the SNP-derived genetic relatedness computed from GCTA using ~544k common variants on HapMap3 (178,075 individual pairs with estimated genetic relatedness ≥ 0.05). The x-axis represents the expected genetic relatedness based on the pedigree information provided by the UKB (monozygotic twin = 1, parent-offspring/full sib = 0.5, second degree relatives = 0.25, third degree relatives = 0.125, and unlabelled pair = 'none') on x-axis. Each circle represents one pair of relatives, the dashed diagonal line represents y = x, and the red horizontal lines represent the mean value of each relatedness group.

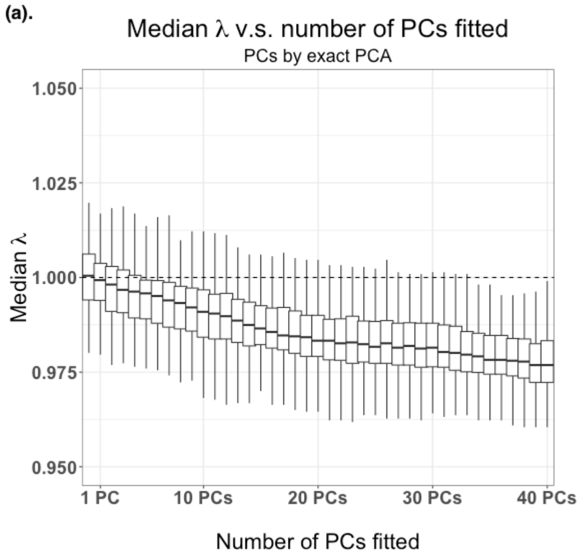


**Supplementary Figure 19.** Relationship between genomic inflation and the number of PCs fitted in LR analysis. The left panel shows the results from analyses fitting PCs computed by PCA using all variants without pruning, and the right panel shows the results from analyses fitting PCs computed by PCA using LD-pruned variants. The phenotypes were adjusted by different number of top PCs (ranging from 1 to 40). The association analyses were performed by LR in PLINK2. The median λ was computed from of all the null variants (i.e., variants on even chromosomes). Each boxplot represents the distribution of median λ across 100 simulation replicates.

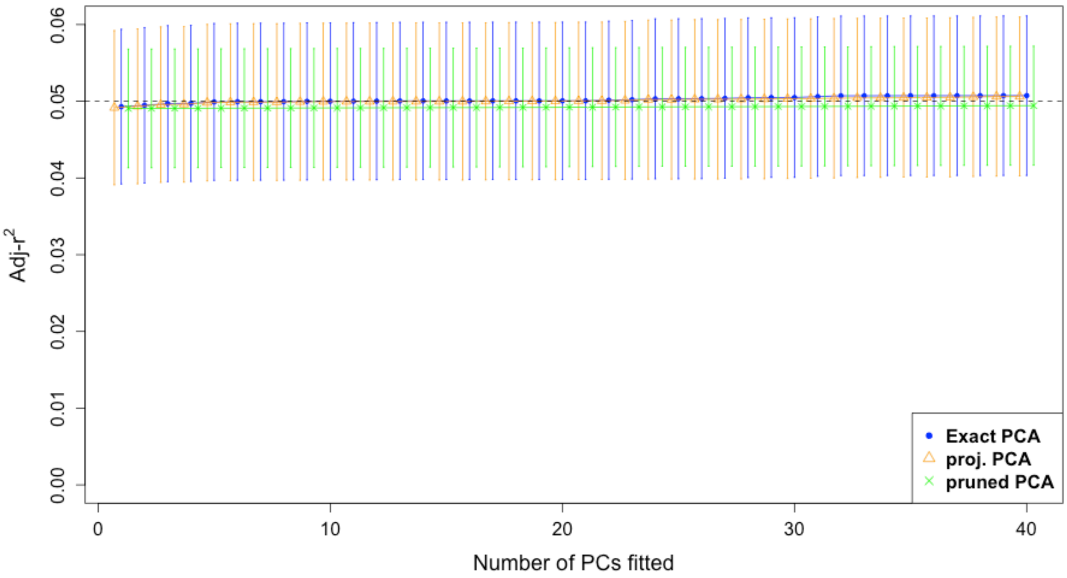


**Supplementary Figure 20.** Proportion of phenotypic variance explained by the top 40 PCs computed from different PCA methods in the simulation. Adjusted $r^2$ (adjusted coefficient of determination) was plotted against the different number of PCs fitted in the model. Three methods, the Exact PCA (Exact PCA, implemented in GCTA) using all variants, the PC projection approach (proj. PCA, implemented in GCTA) using all variants, and flashPCA2 (pruned PCA) using a set of LD-pruned variants, are compared. The dash line represented the parameter used to simulate the proportion of variance explained by population stratification. Each dot represents the average across 100 simulation replicates.
